# rAAV-Delivered Bicistronic Artificial microRNAs for Allele-Specific Silencing Improve Motor and Molecular Outcomes in Spinocerebellar Ataxia Type 3

**DOI:** 10.1101/2025.10.02.680011

**Authors:** Ana Carolina Silva, Carina Henriques, Diana D. Lobo, Ana Rita Fernandes, Miguel M. Lopes, Kevin Leandro, Dina Pereira, Sónia P. Duarte, Sara M. Lopes, Magda M. Santana, Amal Dakka, Steve De Marco, Marla Weetall, Jana Narasimhan, Anu Bhattacharyya, Rui Jorge Nobre, Luís Pereira de Almeida

## Abstract

Spinocerebellar ataxia type 3 (SCA3), also known as Machado-Joseph disease, is an autosomal dominant neurodegenerative disorder caused by the expansion of CAG trinucleotide repeats in the *ATXN3* gene. This mutation induces a toxic gain-of-function of the ATXN3 protein, leading to neurodegeneration, particularly in the cerebellum and brainstem. Despite extensive research, no disease-modifying treatments are available for SCA3 patients.

In this study, we developed and tested a novel therapeutic strategy using recombinant adeno-associated virus (rAAV) to deliver bicistronic artificial microRNAs designed to selectively silence the mutant *ATXN3* allele. Through *in vitro* screening, we identified a lead construct (miATXN3-10x2) that effectively and specifically silenced the mutant allele by targeting of a single nucleotide polymorphism (SNP) associated with the repeat expansion. This construct was packaged into rAAV9 and delivered via intra-cerebellar administration into two mouse models of SCA3, resulting in robust suppression of mutant ATXN3 in the cerebellum. To assess long-term efficacy, we performed intra-cisterna magna (ICM) injections of rAAV9-miATXN3-10x2 in a severe SCA3 transgenic mouse model. Widespread distribution of viral vectors and miATXN3 copies was observed in disease-relevant brain regions. Treated animals exhibited significant and sustained improvements in motor function at 5, 8, and 11 weeks post-injection. Histological analyses showed a reduction in mutant ATXN3 aggregates and a trend toward preventing shrinkage of cerebellar molecular layer. These findings were supported by dose-dependent reductions in mutant *ATXN3* mRNA levels and decreased expression of neuroinflammatory markers in the cerebellum. Additionally, a significant increase of the neuronal marker NeuN was also observed in treated animals. Finally, transcriptomic profiling of the cerebellum demonstrated that treated transgenic animals exhibited an improved transcriptomic signature, shifting toward a wild-type profile.

In conclusion, our findings highlight the therapeutic potential of a single administration of rAAVs encoding bicistronic artificial microRNAs for allele-specific gene silencing in SCA3. This study provides compelling preclinical evidence supporting the translation of this approach into clinical applications for SCA3 patients.

## INTRODUCTION

Expansions of unstable nucleotide repeats are a well-established genetic cause of numerous neurological diseases, with trinucleotide repeat disorders being the most common (reviewed in Paulson, 2018). Spinocerebellar ataxia type 3 (SCA3), or Machado Joseph disease, is the most common form of dominantly inherited ataxia (reviewed in Paulson, 2007; De Mattei et al., 2024), caused by an abnormal expansion of the CAG trinucleotide sequence within the *ATXN3* gene (Kawaguchi et al., 1994; Takiyama et al., 1993).

Intragenic single nucleotide polymorphisms (SNPs) flanking the CAG repeats have been identified, including rs1048755 (A^669^TG/G^669^TG), rs12895357 (C^987^GG/G^987^GG) and rs7158733 (TAA^1118^/TAC^1118^) (Gaspar et al., 2001; Lopes et al., 2020; Elter et al., 2024). Interestingly, specific SNP variants in linkage disequilibrium with the disease-causing mutation were identified and the ACA haplotype was associated with expanded alleles in 72% of SCA3 families worldwide (Gaspar et al., 2001). This knowledge allows further discrimination between the non-expanded and the expanded allele, which can have large implications in the implementation of future gene therapy clinical trials for SCA3.

The CAG expansion in the *ATXN3* gene leads to the production of a mutant ATXN3 protein with an elongated polyglutamine (polyQ) tract. This mutant protein disrupts cellular homeostasis by acquiring toxic properties, including an increased propensity for proteolytic cleavage (Jeannette Hübener et al., 2013; Simões et al., 2012, 2014), misfolding and aggregation (Paulson et al., 1997; Schmidt et al., 1998; Seidel et al., 2010). Several cellular mechanisms have been shown to be compromised in the presence of mutant ATXN3, including the ubiquitin-proteosome system (Burnett et al., 2003; Chai et al., 1999; Donaldson et al., 2003; Holmberg et al., 2004) and autophagy (Ashkenazi et al., 2017; Isabel Onofre et al., 2016; Nascimento-Ferreira et al., 2011; Sittler et al., 2018; Vasconcelos-Ferreira et al., 2021). Other mechanisms have also been implicated in SCA3 pathology, including mitochondrial dysfunction, oxidative stress, transcriptional dysregulation, aberrant protein interactions, and disruptions in axonal transport (reviewed in Nóbrega et al., 2018). Additionally, increasing evidence in polyQ disorders suggests that the expanded mRNA may also contribute to disease pathology (reviewed in Chan, 2014; Griesche et al., 2016) and there is growing evidence that somatic expansion of CAG repeats in the brain may contribute to SCA3 pathogenesis, similar to mechanisms observed in Huntington’s disease (Aldous et al., 2024; Sidky et al., 2024). Overall, the CAG/polyQ expansion in mutant ATXN3 triggers a cascade of pathological processes that ultimately lead to neurodegeneration.

This disorder is characterized by progressive cerebellar ataxia, along with other symptoms such as motor coordination deficits, pyramidal/extrapyramidal signs, oculomotor abnormalities, dysarthria, rigidity, peripheral neuropathy with muscle atrophy, and double vision (Coutinho & Andrade, 1978; Lima & Coutinho, 1980; reviewed in Mendonça et al., 2018). These symptoms arise from progressive neuronal dysfunction and degeneration in the central nervous system (CNS), including the thalamus, midbrain, cerebellum, brainstem, and spinal cord (Koeppen, 2018). The onset of symptoms typically occurs in midlife and CAG repeat length correlates with age of onset variation and rate of progression (Kieling et al., 2007; Diallo et al., 2018; Leotti et al., 2021; Sena et al., 2025). On average, SCA3 patients have a survival period of approximately 20 years after disease onset (Kieling et al., 2007). As of now, no disease-modifying therapies are available for SCA3.

Molecular therapies that induce gene silencing hold significant promise for the treatment of SCA3. Indeed, several studies have demonstrated that silencing of the *ATXN3* gene through antisense oligonucleotide (ASOs) (Moore et al., 2017; McLoughlin et al., 2018, 2023a, 2023b; Bushart et al., 2021; Hauser et al., 2022; Schuster et al., 2024) or RNA interference (RNAi) (Alves et al., 2008, 2010; Rodriguez-Lebron et al., 2013; Costa et al., 2013; Nóbrega et al., 2013, 2014; Conceição et al., 2016; Martier et al., 2019; Nobre & Pereira de Almeida, 2020; Nobre et al., 2022; Rufino-Ramos et al., 2023; Li et al., 2024) have exhibited therapeutic potential in preclinical models. By eliminating ATXN3 transcripts, these approaches may prevent the downstream toxic effects, and thereby slow down or even halt disease progression. Despite their promise, previous preclinical studies are limited in performing a comprehensive phenotypic evaluation, particularly in assessing motor performance. Moreover, they have been based on the use of delivery vectors, silencing molecules and/or invasive administration routes that are hard to translate into clinical settings.

To address these gaps, we sought to optimize and assess the efficacy of recombinant adeno-associated virus serotype 9 (rAAV9)-based gene silencing therapy using artificial miRNAs specifically targeting the mutant *ATXN3* transcript (miATXN3), while also exploring the feasibility of intra-cisterna magna (ICM) injection as a less invasive route of administration for improved clinical application.

Our approach involved the design of novel bicistronic silencing constructs, followed by *in vitro* assessment of allele specificity and gene silencing efficacy. Therapeutic efficacy was next validated in two distinct SCA3 animal models using rAAV9 and two administration routes, intra-parenchymal and cerebrospinal fluid delivery. This treatment resulted in effective transgene delivery, robust and sustained improvement of motor deficits, and amelioration of multiple neuropathological parameters. Moreover, transcriptomic profiling of the cerebellum of treated animals exhibited an improved transcriptomic signature that shifted toward a wild-type profile. Our findings demonstrate that our rAAV-based RNAi gene therapy holds substantial potential as a treatment for SCA3.

## MATERIALS & METHODS

### Silencing construct and rAAV production

Two novel bicistronic constructs, designated miATXN3-8+10 and miATXN3-10x2 (schematic representation in **Figure 1A and B**), were designed based on artificial miRNA sequences previously generated in our laboratory (Nobre & Pereira de Almeida, 2020). The miRNA sequences were designed to target human mutant *ATXN3* mRNA, specifically at SNP variants rs1048755 and/or rs12895357, located in exon 8 and exon 10, respectively. These variants are in linkage disequilibrium with the expanded allele and prevalent in the majority of SCA3 patients (Gaspar et al., 2001). The nucleotide sequences of target *ATXN3* mRNA and the corresponding miRNAs are provided in Supplementary Table 1. Each construct includes two miRNA silencing sequences, both incorporated into a human mir155 scaffold, that were introduced into the same expression cassette, under the control of the CMV early enhancer/chicken β-actin (CAG) promoter.

**Figure 1.**
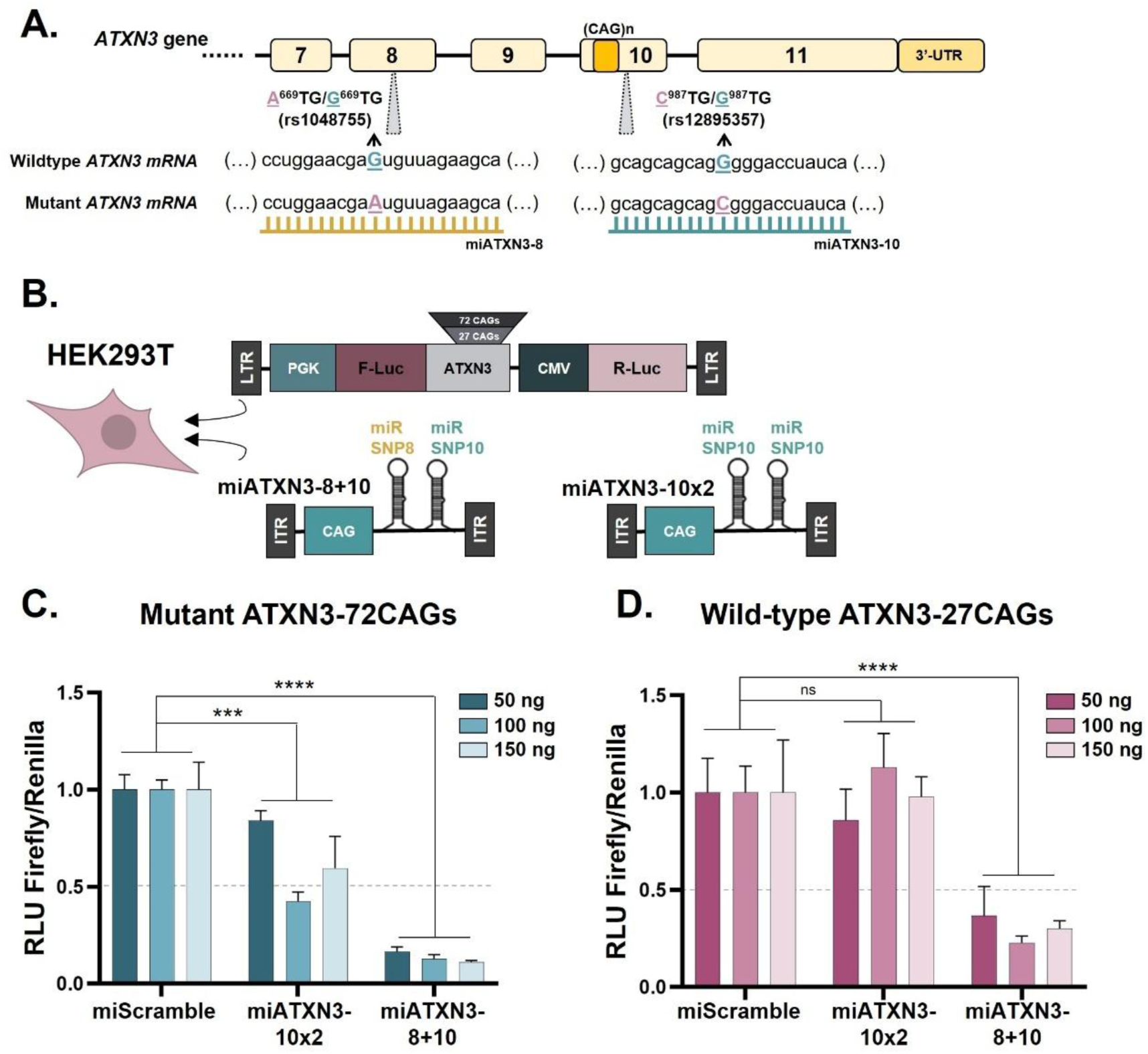
Effective and specific silencing of mutant ATXN3 mRNA by bicistronic artificial microRNAs in HEK293T cells. **(A)** Schematic representation of the *ATXN3* gene with the nucleotide sequences of wild-type and mutant alleles, including the single nucleotide polymorphisms (SNPs) targeted by the artificial microRNAs (miRNAs) directed against SNP8 (miATXN3-8) and SNP10 (miATXN3-10). **(B)** Diagram of the *in vitro* assay in HEK293T cells. Briefly, HEK293T cells were co-transfected with a plasmid encoding either human mutant ATXN3 mRNA (ATXN3-72CAGs) or wild-type ATXN3 mRNA (ATXN3-27CAGs) fused to Firefly luciferase (F-Luc), along with Renilla luciferase (R-Luc) and the novel silencing constructs miATXN3-8+10, miATXN3-10x2 or control scramble. These silencing constructs encode bicistronic artificial miRNAs specifically targeting SNPs associated with the human mutant *ATXN3* allele. Cell lysates were analyzed to evaluate F-Luc and R-Luc luminescence in HEK293T cells expressing ATXN3-72CAGs **(C)** or ATXN3-27CAGs **(D)**. miATXN3-8+10 construct demonstrated robust and efficient silencing of both ATXN3 forms, whereas construct miATXN3-10x2 showed significant silencing of mutant ATXN3 mRNA (ATXN3-72CAGs), while preserving wild-type (ATXN3-27CAGs) expression levels. Data is normalized to luminescence values of miScramble control. The grey dashed line represents 50% silencing. Results are presented as mean±SEM, with n=2-4 replicates. Statistical analysis was performed using Two-way ANOVA with a main effects only model, followed by Dunnett’s multiple comparisons post-hoc test. (ns-not significant).

The expression construct used in the biodistribution assay following ICM injection was previously generated in our laboratory (Nobre & Pereira de Almeida, 2020). This construct contains a single miRNA specifically targeting human mutant *ATXN3*, incorporated into the mir155 scaffold and driven by the U6 promoter. In addition, it carries the enhanced green florescent protein (GFP) sequence under the control of the chicken β-actin (CBA) promoter.

Self-complementary rAAV9 vectors were produced by the CRO PackGene Biotech, LLC (Worcester, MA, USA). Concentrated viral stocks were suspended in sterile 1x Phosphate Buffered Saline (PBS; Fisher BioReagents) supplemented with 0.001% Pluronic F68 (Gibco). Recombinant AAV viral genome (vg) titers (vg/mL) were determined using droplet digital PCR (ddPCR) for specific sequences in AAV2-derived inverted terminal repeats (ITRs). Upon delivery, viral stocks were aliquoted and stored at -80°C until further use.

### In vitro assays

Human embryonic kidney 293 T (HEK293T) cells were cultured in Dulbecco’s Modified Eagle’s Medium (DMEM; Gibco) supplemented with 10% fetal bovine serum (FBS; Biowest) and 1% penicillin/streptomycin (ThermoFisher). Cells were maintained at 37°C in a humidified atmosphere of 5% CO_2_/air. Sterile PBS (pH=7.4) and 0.05% trypsin (Gibco) were used for cell passaging and maintenance.

For *in vitro* assays, 4x10^4^ HEK293T cells were seeded in 48-well plates one day prior to transfection. Twenty-four hours after seeding, cells were co-transfected with a plasmid encoding either human mutant *ATXN3* with 72 CAGs or wild-type *ATXN3* with 27 CAGs fused to Firefly luciferase and Renilla luciferase (Alves et al., 2008; Rufino-Ramos et al., 2023), along with plasmids encoding the two novel bicistronic miRNAs constructs targeting mutant *ATXN3* mRNA. Transfections were carried out using polyethylenimine (PEI; Polysciences) transfection reagent. **Figure 1B** provides a schematic representation of all constructs used.

Forty-eight hours post transfection, the cells were lysed using passive lysis buffer and stored at -80°C. Firefly and Renilla luciferase luminescence signals were subsequently measured using the Dual-Luciferase reporter assay system (Promega), following the manufacturer’s protocol. Firefly and Renilla luciferase activities in each sample were measured using the FLUOstar Omega Microplate Reader (BMG Labtech). Data was normalized to luminescence values of a miScramble control.

### Animal experiments

All experimental procedures with mice were approved by the ethics committee of the Center for Neuroscience and Cell Biology from the University of Coimbra (ORBEA_289_2021/10122021; ORBEA – "Orgão Responsável pelo Bem-Estar Animal”), the Portuguese authority responsible for the regulation of animal experimentation (DGAV 0421/000/000/2022; DGAV – “Direção Geral da Agricultura e Veterinária”) and adhered to the European Community directive (2010/63/EU) for the protection of animals used in scientific purposes. All efforts were made to minimize animal suffering. All researchers responsible for handling animals received proper training (FELASA-certified course) and certification from the Portuguese authorities (DGAV).

### Intraparenchymal cerebellar injections

Intraparenchymal injections were performed by bilateral injection into the deep cerebellar nuclei (DCN). Concentrated viral stocks were thawed on ice and diluted to the desired experimental doses with the vehicle solution. Mice were anesthetized by intraperitoneal administration of a mixture of medetomidine (1 mg/kg, Sedator, Dechra) and ketamine (75 mg/kg, Nimatek, Dechra). A 10 µL Hamilton syringe holding a 30-gauge needle with point style 3 (Hamilton Company) was used.

Ten week old SCA3 YAC84Q transgenic hemizygous mice (Cemal et al., 2002) were used for the first target engagement study. Each mouse received a total of 4 μL of rAAV9-miATXN3-10x2 (2 μL in each brain hemisphere) at three ascending doses (2.4x10^10^ vg n=7; 4.8×10^10^ vg n=7; 9.6×10^10^ vg n=7), or vehicle solution (PBS/0.001% Pluronic F68 solution n=7). Stereotaxic injections in the right and left DCN were performed using the following coordinates relative to bregma: anteroposterior: -6; lateral: -2/+2; ventral: -3.3, at a rate of 0.5 μL/minute.

Five week old SCA3 69Q transgenic mice (Torashima et al., 2008) were used for the second target engagement study. Each mouse received a total of 4 μL of rAAV9-miATXN3-10x2 (2 μL in each brain hemisphere) using two vector doses (9.6×10^9^ vg n=10; 2.4×10^10^ vg n=10), or vehicle solution (PBS/0.001% Pluronic F68 solution n=9). Stereotaxic injections in the right and left DCN were performed using the following coordinates: anteroposterior: -6; lateral: -1.7/+1.7; ventral: -2.6 to -2.7, at a rate of 0.5 μL/minute.

Following surgery, each animal was sutured and given atipamezole (2.5 mg/kg, Antidorm, Calier) and buprenorphine (0.1 mg/kg, Bupaq, Richter Pharma) intraperitoneally. Animals were transferred to a warm cage and kept under supervision until they were fully recovered from anesthesia and then returned to their home cage. The health conditions of all animals were closely monitored during the whole experimental period.

### Intra-cisterna magna injections

Intra-cisterna magna injections were performed using a freehand surgical technique for the biodistribution assay in neonatal SCA3 84Q hemizygous mice (n=7) and for the efficacy study in neonatal SCA3 69Q mice (n=31) and wild-type littermates (n=13). On postnatal day 1 (PND1), pups were carefully placed in a heated pad with shredded nesting material from their dam’s cage. Next, pups were weighed and placed onto aluminum foil on ice for 5-8 minutes to induce hypothermia anesthesia until no movement was observed. Each pup was then held between the index finger and thumb with the head bent to reveal a triangular indentation on the skin near the upper neck area. Injections were performed using a 5 μL Neuros Model Hamilton syringe equipped with a point style 4 and 33 gauge needle (Hamilton Company). A total of 2 μL of rAAV9-miATXN3-10x2 (dose 1: 9.60×10^9^ vg and dose 2: 2.4×10^10^ vg) were delivered into the cisterna magna through the middle of the indentation to a depth of approximately 0.3 mm. Non-treated animals were used as controls. After injection, the needle was left in place for a few seconds to minimize backflow before gentle withdrawal and release of the pup’s head. Pups were placed on a heated pad for recovery and then returned to the dam’s cage. To minimize maternal disturbance, only half of the litter underwent the procedure at a time, regardless of litter size.

### Behavioral tests

All behavioral tests were conducted in the same room after acclimatization, under a 12 hour light/dark cycle, during the animals’ active dark phase. Motor behavior was assessed in all transgenic SCA3 69Q mice by stationary rotarod, beam walking, and Catwalk XT tests at 5, 8 and 11 weeks post-injection.

#### Stationary rotarod

Motor coordination and balance were evaluated using a rotarod apparatus (Panlab, Harvard Apparatus). Mice were placed on the rotarod at a constant speed of 5 rpm for a maximum of 5 minutes, and the latency to fall was recorded. Each mouse performed 3-4 trials over three consecutive days. For analysis, the mean latency to fall from the rotarod was calculated from the three best trials on the second and third days.

#### Beam walking test

Mice were trained to cross an elevated beam to reach an enclosed escape platform. A square beam with 18 mm width was raised to a height of 21 cm with a 40 cm walking distance. Each animal performed two trials of crossing the beam and the latency to cross (up to 60 seconds) was recorded with a camera. Animal performance was then evaluated according to the time to cross the entire beam, which was posteriorly assessed using the Boris software (Version 7.10.5). The mean time of both trials was used for the final values.

#### Catwalk XT

Gait analysis was performed using the CatWalk Automated Gait Analysis System (Noldus Information Technology; version XT 10.6). The runway width was set to 4.1 cm, and the camera height to 35 cm. Camera gain and intensity threshold were set to 19.41 and 0.10, respectively. The home cage of each mouse was placed at the end of the runway to encourage crossings. A minimum of six compliant runs per mouse were acquired. Compliant runs were defined as runs lasting 0.5-15 seconds. After acquisition, each individual compliant run was classified according to pre-defined parameters: minimum number of consecutive steps per run was 12, average speed allowed was 12.5-30 cm/s and maximum allowed speed variation was 30%. Gait parameters analyzed included stride length and paw overlap. Stride length was defined as the average distance between consecutive steps for each paw. The mean distance between each stride for the hind and front left paws, along with hind and front right paws, was calculated. Footprint overlap was quantified as the distance between the center of the hind and front paw footprints.

### Euthanasia and tissue preparation

All animals were euthanized with an overdose of Ketamine (160 mg/kg, Nimatek, Dechra) and Xylazine (8 mg/kg, Sedaxylan, Dechra) and intracardially perfused with cold PBS. The brains were then carefully removed, and the left and right hemispheres were separated. The right hemisphere was dissected to collect different brain regions, then immediately snap-frozen and stored at -80°C for subsequent DNA, RNA and protein extraction. The left hemisphere was post-fixed in 4% paraformaldehyde (Acros Organics) at room temperature for 24 hours, cryoprotected by incubation in 25% sucrose in PBS at 4°C for 48 hours and then stored at -80°C until further processing.

### Histological staining

Sagittal brain sections (30 μm thick) were retrieved from the left hemisphere using a cryostat (CryoStar NX50, Thermo Scientific) at −20°C. These were collected as free-floating sections in 48-well plates containing PBS supplemented with 0.05% sodium azide (Sigma-Aldrich) and stored at 4°C.

For immunofluorescence analysis, sagittal brain slices were selected for visualization of GFP protein and haemagglutinin-tagged (HA) mutant ATXN3 aggregates. Briefly, sections were blocked and permeabilized in a solution of 0.1% Triton X-100 (Fisher Reagents) and 10% normal goat serum (Gibco) in PBS for 1 hour at room temperature under gentle agitation. Next, primary antibody incubation was performed overnight at 4°C using rabbit anti-GFP (1:1000, Invitrogen a6455) or anti-HA (1:1000, Abcam ab9110) prepared in blocking solution. Sections were then washed twice with PBS and incubated for 2 hours at room temperature with a secondary antibody, goat anti-rabbit conjugated to Alexa Fluor 488 (1:250, Invitrogen A11008) or Alexa Fluor 647 (1:250, Invitrogen A21245), prepared in blocking solution. Nuclear staining was performed using 4′,6-diamidino-2-phenylindole (DAPI, Invitrogen). Following three washes with PBS, sections were mounted on gelatin-coated slides, air-dried and coverslipped using Dako fluorescence mounting medium (Agilent).

For histological staining with cresyl violet dye, sagittal brain sections were mounted on gelatin-coated slides and air-dried at room temperature. Then, the slides were immersed in cresyl violet for 5 minutes to stain the Nissl substance in neuronal cell bodies. After staining, sections were washed in ultra-pure H_2_O, followed by dehydration using ethanol solutions (70%, 96%, and 100%; Fisher Chemical). Following a clearing step in xylene substitute (Thermo Scientific), sections were mounted with coverslips using the Richard-Allan Scientific Mounting Medium (Thermo Scientific).

### Quantitative analysis of haemagglutinin-tagged (HA) ATXN3 aggregates

Following HA immunofluorescence staining, images of brain slices were acquired using a Zeiss Axio Scan.Z1 slide scanner microscope (Carl Zeiss Microscopy GmbH), equipped with a 20x objective. Three sagittal sections per animal, corresponding to planes 0.84, 1.08 and 1.32 mm lateral to the midline (according to the Mouse Brain Atlas by Franklin and Paxinos, 2001, 2nd edition), were selected for analysis. The number of HA-tagged ATXN3 aggregates within the cerebellar lobules was manually counted and the lobular area was determined using the QuPath bioimage analysis software (version 0.5.1) (Bankhead et al., 2017). The number of aggregates was normalized to the respective lobular area. Final values were expressed as the average number of aggregates per square millimeter (aggregates/mm^2^) across the three selected sections for each animal.

### Quantitative analysis of cerebellar layers

Following cresyl violet staining, images of the whole cerebellum were acquired using the Zeiss Axio Scan.Z1 slide scanner microscope with a 20x objective and analyzed using the Zen Blue software (blue edition, Version 3.8, Carl Zeiss Microscopy GmbH). For each animal, three sagittal brain sections (0.84, 1.08 and 1.32 mm lateral to the midline according to the Mouse Brain Atlas by Franklin and Paxinos, 2001, 2nd edition) were selected to measure the thickness of molecular layer in the cerebellar lobules.

For each lobule within a section, molecular layer thickness was manually calculated by taking three independent measurements at predefined regions. The final thickness value for each value was determined as the mean of these three measurements. The overall molecular layer thickness for each animal was calculated as the average of all lobules across the three selected sections.

### Total DNA, RNA and protein purification

Simultaneous purification of genomic DNA, total RNA, and total protein from the same tissue sample was performed using the AllPrep DNA/RNA/Protein kit (Qiagen). Briefly, tissues were disrupted using a pestle and homogenized with a syringe with 21 gauge needle, in 600 µL of lysis buffer. The remaining procedure was performed according to the manufacturer’s instructions. RNA and DNA concentrations were calculated using the Nanodrop 2000 Spectrophotometer (Thermo Scientific) and sample purity was assessed by determining the optical density (OD) ratio at 260/280 nm. The protein pellet was resuspended in a solution of 8M Urea and 1% sodium dodecyl sulfate (SDS) in 100mM Tris-HCl (pH 8.0), followed by sonication with 2 series of 3-4 seconds ultra-sound pulses (1 pulse/sec). RNA and protein extracts were stored at -80°C, while DNA samples were stored at -20°C.

### Quantification of rAAV genome copy number by real-time quantitative PCR

Recombinant AAV genome copy number was determined using the AAVpro™ Titration Kit (Takara Bio Inc). Genomic DNA was diluted to a concentration of 10 ng/μL and used as a template for real-time quantitative PCR (qPCR). For each reaction, 50 ng of total DNA was analyzed. The AAV2-derived ITR sequence was targeted to quantify the number of viral genomes present in each sample. qPCR was performed using the Applied Biosystems StepOnePlus Real-Time PCR system (Life Technologies). Cycle threshold values were determined using the StepOnePlus software (version 2.3, Life Technologies). Viral genome concentration (vg/μL) was calculated from a standard curve generated from serial dilutions of a positive control provided by the kit. Final values of viral genomes were normalized and expressed as vg/μg of total DNA.

### DNase treatment, cDNA synthesis and Reverse Transcription qPCR

RNA samples underwent DNase digestion using the RNase-Free DNase Set (Qiagen), to eliminate genomic DNA contamination and prevent co-amplification. Subsequently, reverse transcription was performed on 500 ng of total RNA, using the iScript cDNA Synthesis Kit (Bio-Rad) and the Veriti Thermal Cycler PCR system (Life Technologies). The resulting cDNA was diluted to a final concentration of 2.5 ng/μl in nuclease-free water. Reverse Transcription qPCR (RT-qPCR) reactions were performed on 10 ng of diluted cDNA using the SsoAdvanced SYBR Green Supermix (Bio-Rad), following the manufacturer’s protocol. Previously validated primers for human *ATXN3*, mouse *Atxn3*, mouse *Hprt* (hypoxanthine guanine phosphoribosyl transferase), mouse Iba1 (Allograft inflammatory factor 1 *Aif1*, also known as ionized calcium-binding adapter molecule 1 Iba1) and mouse *Gfap* (Glial fibrillary acidic protein) were used for mRNA expression analysis. The sequences of all primers used in RT-qPCR are listed in Supplementary Table 2. Appropriate negative controls were included in all experiments. Each reaction was performed in duplicate under the following cycling conditions: 95°C for 30 seconds (initial denaturation), followed by 45 cycles at 95°C for 5 seconds (denaturation), and 56°C for 15 seconds (annealing/extension). The protocol was performed using the CFX96 Real-Time PCR system (Bio-Rad) and the Cycle threshold values were determined using the Bio-Rad CFX Maestro software (version 1.0). Internal controls for normalization of RT-qPCR data were performed using mouse *Hprt* or *Atxn3* mRNA levels. Relative gene expression between control and experimental samples was calculated through the 2^-ΔΔCt^ method.

### MicroRNA Taqman assays

Reverse transcription of RNA samples was performed using the TaqMan MicroRNA Reverse Transcription Kit (Applied Biosystems), according to the manufacturer’s instructions. Briefly, total RNA was diluted to a concentration of 1 ng/μL, with 5 ng of RNA used as the template for each reverse transcription reaction. Reverse transcription was carried out using the Veriti Thermal Cycler. A custom TaqMan small RNA assay (Applied Biosystems) was used to design microRNA Taqman probes specifically for the detection of miATXN3-10x2. All TaqMan small RNA reactions were performed according to the manufacturer’s protocol on StepOnePlus Real-Time PCR system. To determine absolute miRNA expression levels, a standard curve was prepared from a commercially synthesized RNA oligonucleotide (Thermo Scientific). Final miRNA levels were normalized to non-treated controls and are presented as miRNA copies per μg of total RNA.

### Western Blotting

Protein concentration was determined using the BCA method (Pierce BCA Protein Assay Kit, Thermo Scientific). Protein samples (90 μg or 60 μg per lane) were mixed with sample buffer (0.5 M Tris-HCl, pH 6.8; 10% SDS; 30% glycerol; 0.6 M DTT; bromophenol blue), vortexed, and denatured at 95°C for 5 minutes.

Proteins were resolved in SDS polyacrylamide gels (SDS-PAGE) composed of 4% stacking gel and 10% running gel. Electrophoresis was conducted at 80 V for 15 minutes, followed by 90 V for approximately 90 minutes. Proteins were transferred to polyvinylidene difluoride (PVDF) membranes (Merck, Millipore) at 400mA for 3 hours in CAPS buffer (Fisher Bioreagents) containing 0.5% methanol (Fisher Chemical) at 4°C, using the Criterion Blotter with Plate Electrodes system (Bio-Rad). Total protein staining was performed in each membrane using the No-Stain Protein Labeling Reagent (Invitrogen), following manufacturer’s instructions and visualized using the ChemiDoc Imaging System (Bio-Rad). Next, membranes were blocked with 5% non-fat milk powder in 0.1% Tween 20 in Tris buffered saline (TBS-T, Fisher BioReagents) for 1 hour at room temperature and incubated overnight at 4°C with primary antibodies: mouse anti-1H9 (1:5000, Millipore MAB5360), mouse anti-β-Actin (1:5000, Sigma A5316), rabbit anti-NeuN (1:1000, Millipore, ABN78), and mouse anti-Gapdh (1:3000, Millipore MAB374), diluted in blocking solution. Membranes were subsequently washed three times with TBS-T and incubated at room temperature for 2 hours with an alkaline phosphatase-conjugated goat anti-mouse secondary antibody (1:10000, Invitrogen 31328) or goat anti-rabbit (1:10000, Invitrogen 31340) prepared in blocking solution. Finally, membranes were washed three times with TBS-T and incubated with Enhanced Chemifluorescent substrate (GE Healthcare). Chemifluorescence signals were visualized using the ChemiDoc Imaging System (Bio-Rad) and quantitative analysis was conducted on the scanned membranes using Image J software (version 1.54f, NIH, USA). Densiometric quantification was normalized to β-Actin or Gapdh protein levels of the corresponding lanes.

### RNA sequencing analysis

Total RNA samples extracted from the cerebellum at 13 weeks post rAAV9-miATXN3-10x2 administration were used to perform transcriptome sequencing. A total of 14 animals from the long-term efficacy study were selected for RNA sequencing analysis, comprising of non-treated SCA3 69Q mice (n=5, 2 males and 3 females), treated SCA3 69Q mice (n=6, 1 male and 5 females), as well as non-treated wild-type littermates (3 females). For this analysis, treated transgenic animals showing more than 30% silencing of human mutant *ATXN3* mRNA in the cerebellum were selected and designated “top treated group”.

Library preparation and RNA sequencing was performed by Novogene (Munich, Germany). Briefly, mRNA was purified from total RNA using poly-T oligo-attached magnetic beads. Following mRNA fragmentation, the first strand of cDNA was synthesized using random hexamer primers, followed by the second cDNA strand synthesis using either dUTP for directional library or dTTP for non-directional library. cDNA libraries were sequenced using Illumina platforms.

For bioinformatic analysis, raw data (raw reads) in FASTQ format was processed using in-house perl scripts. Clean data (clean reads) was acquired by removing reads containing adapter and ploy-N, and low quality reads from raw data. Next, reads were mapped to the reference genome “ensembl_111_mus_musculus_grcm39_primary” and paired-end clean reads were aligned to the reference genome using Hisat2 (v2.0.5). The number of read counts mapped to individual genes were calculated using FeatureCounts (v1.5.0-p3)

Differential expression analysis was done using the NovoMagic Online RNA-seq Bioinformatics Analysis Tool (Novogene) with the DESeq2 R package (1.20.0). P values were adjusted using the Benjamini-Hochberg method to control the false discovery rate. Genes with an adjusted p value < 0.05 found by DESeq2 and |log2FoldChange| > 0 were defined as differentially expressed.

Gene Ontology (GO) and Kyoto Encyclopedia of Genes and Genomes (KEGG) enrichment analyses of differentially expressed genes (DEGs) were executed by the R (Version 3.0.3) clusterProfiler package following DEG analysis and annotation to the GO and KEGG database, respectively. GO terms and KEGG pathways with an adjusted p value < 0.05 were considered significantly enriched for all DEG sets.

Principal component analysis (PCA) was performed on the normalized read count data of the different experimental groups using the R (Version 3.0.3) ggplot2 package.

### Statistical analysis

Statistical analysis was performed using the GraphPad Prism software (version 9.0.0). All data are presented as mean ± standard error of the mean (sem). Correlation analysis was conducted using Person’s correlation test. Outliers were removed according to the Grubbs test (α=0.05). Two way ANOVA, One way ANOVA followed by Dunnett’s test or Student’s t-test were conducted when appropriate. Statistical significance thresholds were defined as follows: * p < 0.05, ** p < 0.01, *** p < 0.001, and **** p < 0.0001.

## RESULTS

### Design and *in vitro* testing of optimized artificial miRNAs targeting mutant *ATXN3* mRNA (miATXN3)

Genetic population studies have identified a higher prevalence of certain haplotypes of intragenic SNPs in the expanded allele of SCA3 individuals (Gaspar et al., 2001; Elter et al., 2024). Allele-specific silencing of mutant *ATXN3* can be achieved through targeting of SNP variants that are in linkage disequilibrium with the CAG trinucleotide repeat expansion in the *ATXN3* mutant allele (**Figure 1A**). Previously, we designed artificial miRNA sequences targeting SNP8 (A^669^TG/G^669^TG: rs1048755), located in exon 8, and SNP10 (C^987^GG/G^987^GG: rs12895357), located in exon 10 (Nobre & Pereira de Almeida, 2020). We specifically targeted the nucleotides A and C, of SNP8 and SNP10, respectively, which are more commonly associated with the expanded *ATXN3* allele (Gaspar et al., 2001). In the present study, we developed two novel allele-specific silencing constructs targeting mutant *ATXN3*, building on these earlier sequences. Each construct included two silencing sequences within a single expression cassette (bicistronic miRNA constructs), with artificial miRNA expression driven by the ubiquitous CAG promoter. One cassette combined an artificial miRNA targeting SNP8 with another targeting SNP10 (miATXN3-8+10), while the second cassette contained two artificial miRNA both directed against SNP10 in the mutant *ATXN3* mRNA (miATXN3-10x2) (**Figure 1B**).

To test the allele specificity and silencing efficacy of these constructs, we used a luminescence reporter system in which human mutant ATXN3 (ATXN3-72CAGs) or wild-type ATXN3 (ATXN3-27CAGs) is fused to Firefly luciferase under the control of the PGK promotor (Alves et al., 2008; Rufino-Ramos et al., 2023). Renilla luciferase, regulated by the CMV promoter, served as normalization control. These constructs encoding ATXN3-72CAGs and ATXN3-27CAGs reporters were then co-transfected in HEK293T cells with either the silencing construct miATXN3-8+10 or miATXN3-10x2 (**Figure 1B**). In cells expressing *ATXN3*-72CAGs, both constructs demonstrated significant silencing of mutant *ATXN3*, with miATXN3-8+10 and miATXN3-10x2 achieving knockdown efficiencies of up to 90% and 58%, respectively (**Figure 1C**). In contrast, in cells expressing *ATXN3*-27CAGs, miATXN3-10x2 did not reduce wild-type *ATXN3* levels, whereas miATXN3-8+10 achieved robust silencing of wild-type mRNA, with up to 77% reduction (**Figure 1D**).

These findings indicate that although miATXN3-8+10 provided strong and efficient silencing of both mutant and wild-type *ATXN3* mRNAs, miATXN3-10x2 preferentially suppressed mutant *ATXN3* while preserving the wild-type form. Based on these results, miATXN3-10x2 was selected as the lead candidate for allele-specific silencing of mutant *ATXN3* mRNA.

### *In vivo* assessment of rAAV9-miATXN3-10x2 as a therapeutic strategy for SCA3: efficient transduction and target engagement following intra-cerebellar administration

Having validated the allele-specific silencing capacity of miATXN3-10x2 *in vitro*, we aimed to assess its therapeutic potential *in vivo* using two different animal models. We first employed the hemizygous SCA3 YAC84Q mouse model (Cemal et al., 2002), which carries two copies of the full-length human mutant *ATXN3* gene, including SNP10 (rs12895357) with the C nucleotide specifically targeted by miATXN3-10x2. To evaluate the distribution and target engagement of our lead therapeutic candidate, we performed intra-cerebellar injections of three escalating vector doses of rAAV9 encoding miATXN3-10x2 at 10 weeks of age (**Figure 2A**).

**Figure 2.**
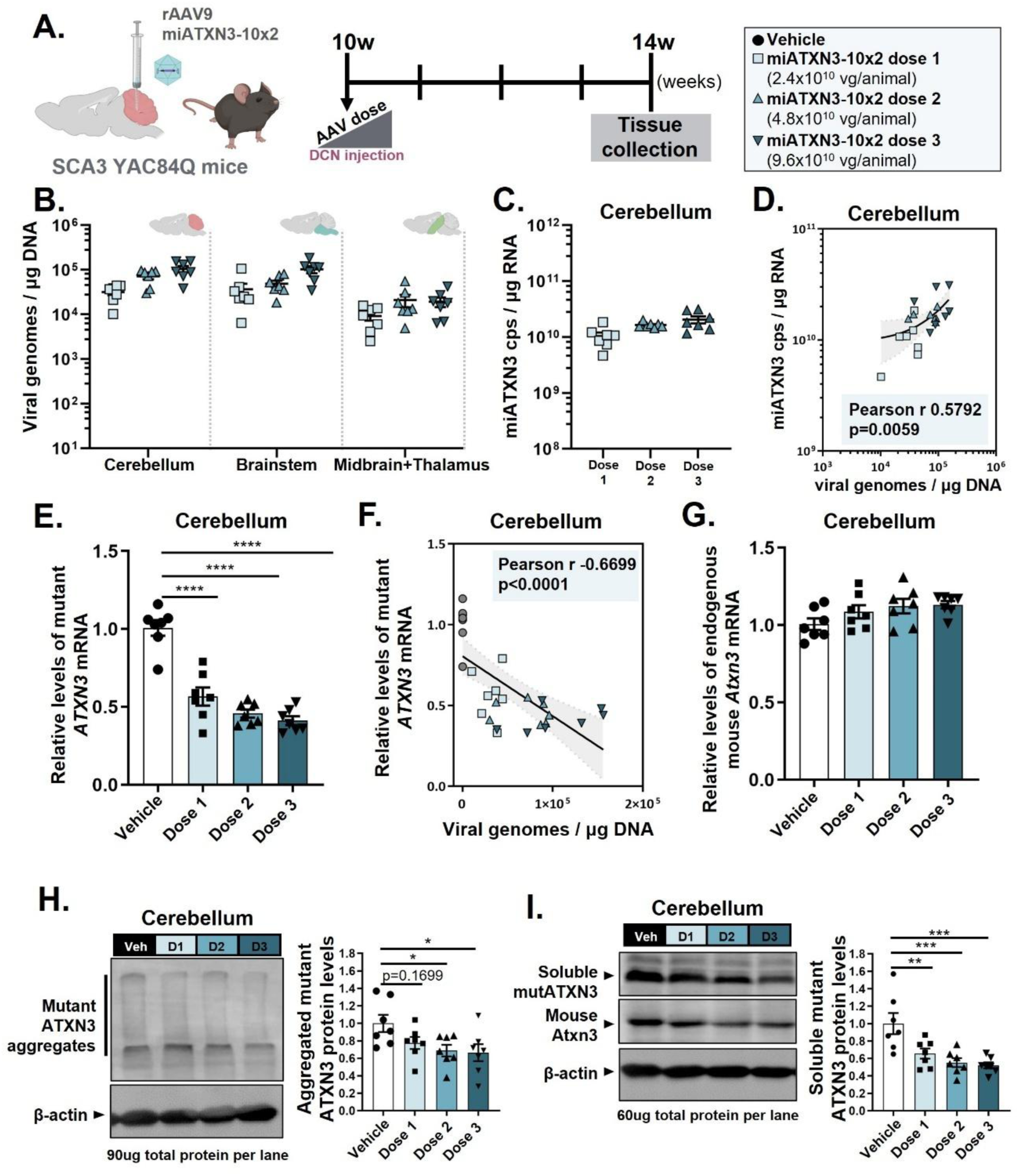
Dose-dependent suppression of mutant ATXN3 following rAAV9-miATXN3-10x2 intra-cerebellar administration in hemizygous SCA3 YAC84Q transgenic mice. **(A)** Schematic representation of the experimental design and timeline. A dose scaling study was conducted using three different doses of rAAV9 packaged with the miATXN3-10x2 silencing construct: dose 1 - 2.4×10^10^ vg/animal; dose 2 - 4.8×10^10^ vg/animal; dose 3 - 9.6×10^10^ vg/animal. At 10 weeks of age, hemizygousSCA3 YAC84Q mice were bilaterally injected in the deep cerebellar nuclei (DCN) with either vehicle or rAAV9-miATXN3-10x2. Four weeks post-surgery, mice were euthanized for brain tissue collection. **(B)** Total DNA was isolated from different brain tissues and qPCR was performed to determine rAAV genome copy number distribution in the cerebellum, brainstem and midbrain+thalamus. No vector genome copies were detected in untreated mice. **(C)** Total RNA was isolated from cerebellar tissue for small RNA Taqman-probe assay. RT-qPCR revealed a dose-dependent increase in the number of miATXN3 copies, with the highest number obtained in the high dose group. **(D)** A positive correlation was observed between viral genomes and miATXN3 copies in the cerebellum of treated mice. **(E)** RT-qPCR analysis of cerebellar tissue showed a robust, dose-dependent reduction in mutant ATXN3 mRNA levels. **(F)** Higher viral genome copy numbers were strongly correlated with lower levels of mutant ATXN3 mRNA in the cerebellum. **(G)** No significant differences were observed in endogenous mouse Atxn3 mRNA levels across all treatment groups. **(H, I)** Representative images of western blot analysis are shown. Quantification of aggregated mutant ATXN3 (H) and soluble mutant ATXN3 (I) protein levels revealed strong, dose-dependent suppression of both protein forms in the cerebellum of treated mice. N=7 mice per experimental group. Data presented as mean ± SEM. Statistical analysis was performed using One-way ANOVA, followed by Dunnett’s multiple comparisons post-hoc test, and Pearson’s correlation test (two-tailed p value; grey area denotes the 95% confidence interval).

Although treated animals showed slightly lower average body weights than vehicle-treated controls in the weeks following treatment, no statistically significant differences were observed between groups at any evaluated time point (**Supplementary Figure 1A**).

Analysis of viral genome distribution confirmed a dose-dependent increase in genome copies in the SCA3 mouse brain, including major affected brain regions in SCA3, i.e. cerebellum and brainstem (Cerebellum - miATXN3-10x2 dose 1: 3.2×10^4^ vg/µg DNA ± 4.8×10^3^, miATXN3-10x2 dose 2: 7.2×10^4^ vg/µg DNA ± 1.0×10^4^ and miATXN3-10x2 dose 3: 1.0×10^5^ vg/µg DNA ± 1.7×10^4^; Brainstem - miATXN3-10x2 dose 1: 3.7×10^4^ vg/µg DNA ± 1.2×10^4^, miATXN3-10x2 dose 2: 4.9×10^4^ vg/µg DNA ± 8.3×10^3^ and miATXN3-10x2 dose 3: 1.0×10^5^ vg/µg DNA ± 1.8×10^4^; Midbrain+Thalamus - miATXN3-10x2 dose 1: 9.2×10^3^ vg/µg DNA ± 1.9×10^3^, miATXN3-10x2 dose 2: 2.1×10^4^ vg/µg DNA ± 6.5×10^3^ and miATXN3-10x2 dose 3: 1.9×10^4^ vg/µg DNA ± 4.7×10^3^) (**Figure 2B**). Similarly, miATXN3 expression was detected in the cerebellum, with a clear dose-dependent increase (Cerebellum - miATXN3- 10x2 dose 1: 1.0×10^10^ cps/µg RNA ± 1.6×10^9^, miATXN3-10x2 dose 2: 1.6×10^10^ cps/µg RNA ± 6.9×10^8^ and miATXN3-10x2 dose 3: 2.0×10^10^ cps/µg RNA ± 2.9×10^9^) (**Figure 2C**). A positive correlation between viral genomes and miATXN3 copy numbers was confirmed in the cerebellum (Pearson r=0.5792, p=0.0059) (**Figure 2D**).

We next evaluated target engagement in disease-relevant brain regions. In the cerebellum, a strong and significant knockdown of *ATXN3* mRNA was observed across all tested doses, with the highest dose achieving up to 60% silencing (vehicle: 1.00 ± 0.05, miATXN3-10x2 dose 1: 0.57 ± 0.06, miATXN3-10x2 dose 2: 0.46 ± 0.03 and miATXN3-10x2 dose 3: 0.41 ± 0.03) (**Figure 2E**). In the brainstem, a significant reduction in mutant *ATXN3* mRNA levels was also observed for the medium and high doses, with silencing efficiencies of up to 30% (vehicle: 1.00 ± 0.04, miATXN3-10x2 dose 1: 0.88 ± 0.06, miATXN3-10x2 dose 2: 0.81 ± 0.04 and miATXN3-10x2 dose 3: 0.70 ± 0.04) (**Supplementary Figure 1B**). Strong negative correlations were identified between viral genome copies and *ATXN3* mRNA levels in both the cerebellum (Pearson r=-0.6699, p<0.0001) (**Figure 2F**) and the brainstem (Pearson r=- 0.6251, p=0.0004) (**Supplementary Figure 1C**). Interestingly, a strong positive correlation was found between mutant *ATXN3* mRNA levels in the cerebellum and the brainstem, with lower levels of mutant *ATXN3* mRNA in the cerebellum correlating with lower transcript levels in the brainstem (Pearson r=0.7987, p<0.0001) (**Supplementary Figure 1D**). Endogenous mouse *Atxn3* mRNA expression remained unchanged across treatment groups in the cerebellum (**Figure 2G**) and the brainstem (**Supplementary Figure 1E**).

Consistent with the high levels of *ATXN3* mRNA silencing in the cerebellum, we observed a dose-dependent reduction of up to 34% in aggregated mutant ATXN3 (vehicle: 1.00 ± 0.10, miATXN3-10x2 dose 1: 0.78 ± 0.07, miATXN3-10x2 dose 2: 0.69 ± 0.07 and miATXN3-10x2 dose 3: 0.66 ± 0.09) (**Figure 2H**). Soluble mutant ATXN3 protein levels were also reduced in the cerebellum, with a maximum reduction of 48% in the group receiving the highest vector dose (vehicle: 1.00 ± 0.12, miATXN3-10x2 dose 1: 0.65 ± 0.06, miATXN3-10x2 dose 2: 0.55 ± 0.05 and miATXN3-10x2 dose 3: 0.52 ± 0.03) (**Figure 2I**). No significant reductions in either protein forms were detected in the brainstem (**Supplementary Figure 1F**). Accordingly, a strong correlation was observed between mutant *ATXN3* mRNA and soluble ATXN3 protein levels in the cerebellum (Pearson r=0.7086, p<0.0001), but not in the brainstem (Pearson r=0.1055, p=0.5932) (**Supplementary Figure 1G**).

We further tested rAAV9-miATXN3-10x2 in a second transgenic SCA3 mouse model, which expresses a truncated human mutant ATXN3 protein with 69Q, exhibiting severe neuropathological alterations from 21 days of age (Torashima et al., 2008). Importantly, this animal model also contains the SNP10 sequence with the C nucleotide. Animals received bilateral intra-cerebellar injections at five weeks of age with two vector doses and were euthanized at 18 weeks to assess vector distribution and target engagement (**Supplementary Figure 2A**). Body weight measurements revealed no significant differences between treated and vehicle groups (**Supplementary Figure 2B**).

We categorized the animals based on the detected number of viral genomes in the targeted brain region (i.e. cerebellum), which resulted in the separation into “low vg/μg DNA” and “high vg/μg DNA” groups (miATXN3-10x2 low vg/μg DNA: 4.3×10^2^ ± 6.8×10^1^ and miATXN3- 10x2 high vg/μg DNA: 3.4×10^4^ ± 7.7×10^3^) (**Supplementary Figure 2C**). Moreover, sustained transgene expression was detected in the cerebellum, emphasized by the high levels of miATXN3 copies in this brain region (miATXN3-10x2 copies in low vg/μg DNA: 1.1×10^8^ ± 2.2×10^7^ and miATXN3-10x2 copies in high vg/μg DNA: 1.6×10^9^ ± 3.2×10^8^) (**Supplementary Figure 2D**). A strong positive correlation was identified between the number of viral genomes and miATXN3 copies in the cerebellum (Pearson r=0.7646, p=0.0001) (**Supplementary Figure 2E**). The high levels of miATXN3 expression were accompanied by a reduction in the levels of mutant *ATXN3* mRNA in the cerebellum of treated animals, with a significant 28% reduction of mutant *ATXN3* mRNA levels in the high vg/μg DNA group (vehicle: 1.00 ± 0.04, miATXN3-10x2 low vg/μg DNA: 0.89 ± 0.05 and miATXN3-10x2 high vg/μg DNA: 0.72 ± 0.05) (**Supplementary Figure 2F**). Of note, higher number of viral genomes were correlated with higher silencing levels of *ATXN3* mRNA (Pearson r=-0.6360, p=0.0002) (**Supplementary Figure 2G**).

In summary, intra-cerebellar administration of rAAV9-miATXN3-10x2 in both SCA3 YAC84Q and SCA3 69Q mice resulted in efficient transduction, sustained miATXN3 expression, and robust suppression of mutant *ATXN3* in the cerebellum. Together, these findings highlight the feasibility of this therapeutic strategy for targeting disease-affected brain regions *in vivo*.

### Widespread and sustained transgene expression of rAAV9-miATXN3-10x2 in SCA3 mouse models following ICM administration

Having established that rAAV9-miATXN3-10x2 can silence mutant *ATXN3* mRNA in SCA3 mouse models following intraparenchymal administration, we aimed to investigate a less invasive delivery method. To this end, we conducted a biodistribution study to evaluate transgene expression throughout the mouse brain following intra-CSF injection in neonatal SCA3 YAC84Q mice (**Figure 3A**). We used a rAAV9 vector encoding GFP as a reporter together with the artificial miRNA sequence targeting mutant ATXN3 mRNA. The vector was delivered via ICM injection at PND1. Fifteen days post-injection, the animals were euthanized, and immunohistochemistry was performed to assess transgene distribution.

**Figure 3.**
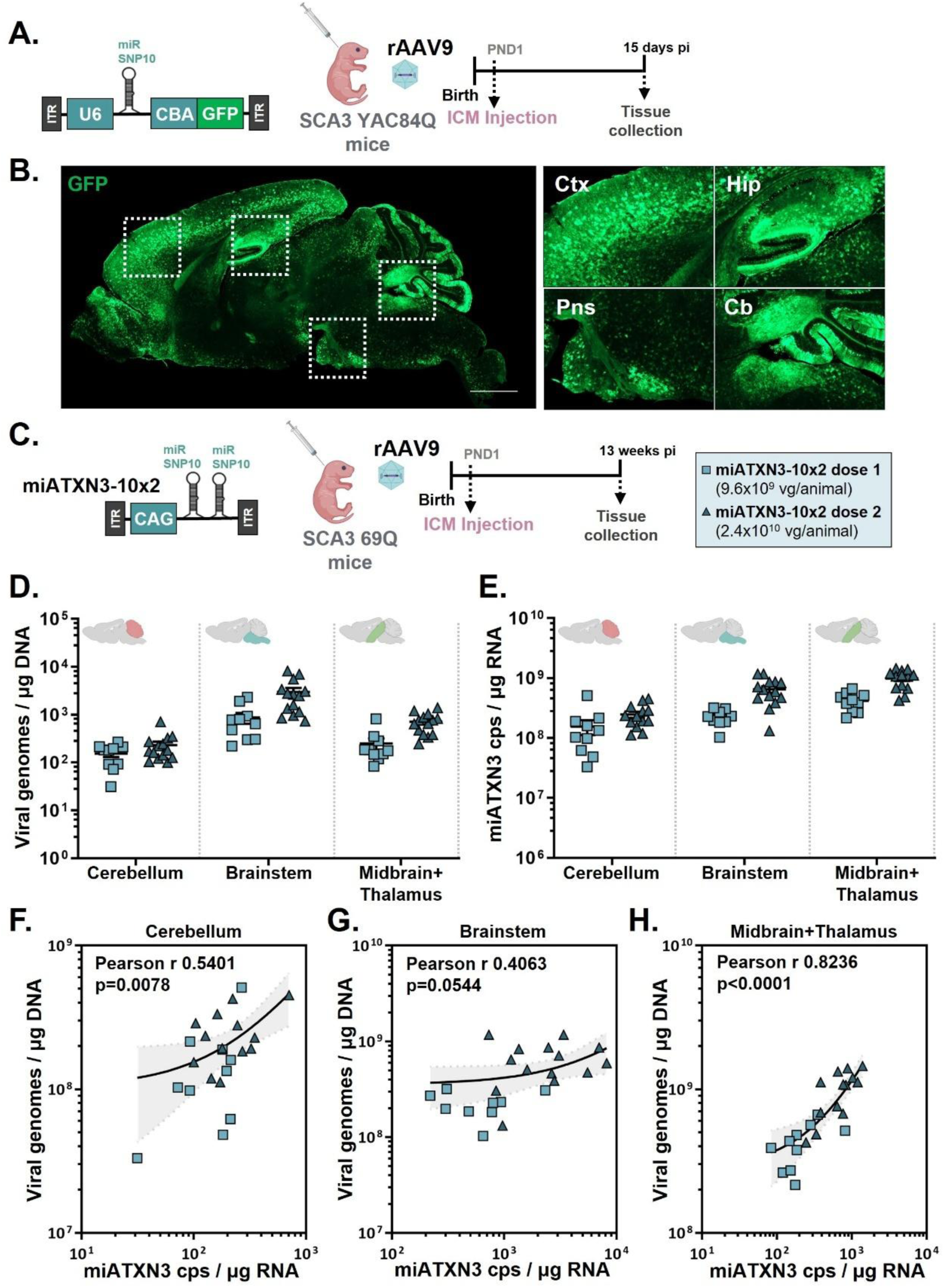
Intra-cisterna magna injection in neonatal SCA3 mice results in widespread rAAV9 transduction throughout the brain. **(A)** Biodistribution assay to evaluate transgene expression across the mouse brain following intra-cisterna magna (ICM) delivery. Hemizygous SCA3 YAC84Q mice received rAAV9 vectors encoding the reporter gene GFP via ICM injection at postnatal day 1 (PND1). **(B)** Immunofluorescent analysis using anti-GFP staining revealed widespread transduction of the brain, including disease-relevant brain regions such as the DCN and pons. Qualitative analysis was performed following immunofluorescence. A representative image of GFP biodistribution is shown. Ctx - cortex; Hip - hippocampus; Pns - pons; Cb – Cerebellum. **(C)** rAAV9- miATXN3-10x2 was delivered via ICM injection at PND1 in a transgenic SCA3 69Q mouse model expressing a truncated mutant ATXN3. Thirteen weeks post-injection, brains were collected to assess viral genome distribution and miRNA expression. **(D)** Quantification of viral genome copies in different brain regions revealed successful transduction. **(E)** Measurement of miATXN3 copies in the cerebellum, brainstem, and midbrain+thalamus confirmed efficient miRNA expression. **(F-H)** Pearson’s correlation analysis showed a significant positive correlation between the number of viral genomes and miATXN3 copies in the cerebellum **(F)**, brainstem **(G)** and midbrain+thalamus **(H)**. N=7-14 mice per experimental group. Data presented as mean ± SEM. Statistical analysis was performed using the Pearson’s correlation test: two-tailed p value; grey area denotes the 95% confidence interval.

Widespread GFP expression was observed throughout the brain, including in key regions affected in SCA3, such as the cerebellum (including the DCN and cerebellar lobules) and the brainstem (pons) (**Figure 3B**).

Next, we conducted a long-term efficacy study by performing ICM injections with two vector doses at PND1 into SCA3 69Q mice and assessed different parameters at 13 weeks post-administration. We began by assessing transduction efficiency through viral genome distribution and miATXN3 copy numbers analysis (**Figure 3C**). Viral genomes were detected in all examined brain regions, with levels varying by dose (Cerebellum - miATXN3-10x2 dose 1: 1.5×10^2^ vg/ug DNA ± 2.4×10^1^ and miATXN3-10x2 dose 2: 2.3×10^2^ vg/ug DNA ± 4.2×10^1^; Brainstem - miATXN3-10x2 dose 1: 8.7×10^2^ vg/ug DNA ± 2.2×10^2^ and miATXN3-10x2 dose 2: 3.0×10^3^ vg/ug DNA ± 6.3×10^2^; Midbrain+Thalamus - miATXN3-10x2 dose 1: 2.5×10^2^ vg/ug DNA ± 6.7×10^1^ and miATXN3-10x2 dose 2: 7.1×10^2^ vg/ug DNA ± 9.0×10^1^) (**Figure 3D**). Similarly, miATXN3 copy number was detected throughout the brain, showing a dose-dependent increase (Cerebellum - miATXN3-10x2 dose 1: 1.6×10^8^ cps/ug RNA ± 4.4×10^7^ and miATXN3-10x2 dose 2: 2.5×10^8^ cps/ug RNA ± 3.0×10^7^; Brainstem - miATXN3-10x2 dose 1: 2.3×10^8^ cps/ug RNA ± 2.3×10^7^ and miATXN3-10x2 dose 2: 6.5×10^8^ cps/ug RNA ± 8.2×10^7^; Midbrain+Thalamus - miATXN3-10x2 dose 1: 4.2×10^8^ cps/ug RNA ± 4.5×10^7^ and miATXN3-10x2 dose 2: 9.9×10^8^ cps/ug RNA ± 9.5×10^7^) (**Figure 3E**).

Further analysis revealed that higher numbers of viral genomes correlated with higher number of miATXN3 copies in the cerebellum (Pearson r=0.5401, p=0.0078) (**Figure 3F**), brainstem (Pearson r=0.4063, p=0.0544) (**Figure 3G**) and midbrain+thalamus (Pearson r= 0.8236, p<0.0001) (**Figure 3H**).

Altogether, we observed widespread and sustained transgene expression following ICM delivery of our therapeutic strategy in the SCA3 mouse brain, suggesting that this administration route holds high promise for rAAV-based gene therapies in neurodegenerative disorders.

### rAAV9-miATXN3-10x2 improves molecular parameters and ameliorates the severe motor phenotype in the SCA3 69Q mouse model following CSF-directed administration

Next, we evaluated the therapeutic potential of rAAV9-miATXN3-10x2 in ameliorating the severe motor deficits and associated molecular alterations that characterize the SCA3 69Q mouse model (Torashima et al., 2008). In the long-term efficacy study, motor performance was evaluated at multiple timepoints, and animals were euthanized 13 weeks post-injection for analysis of relevant biochemical endpoints (**Figure 4A**).

**Figure 4.**
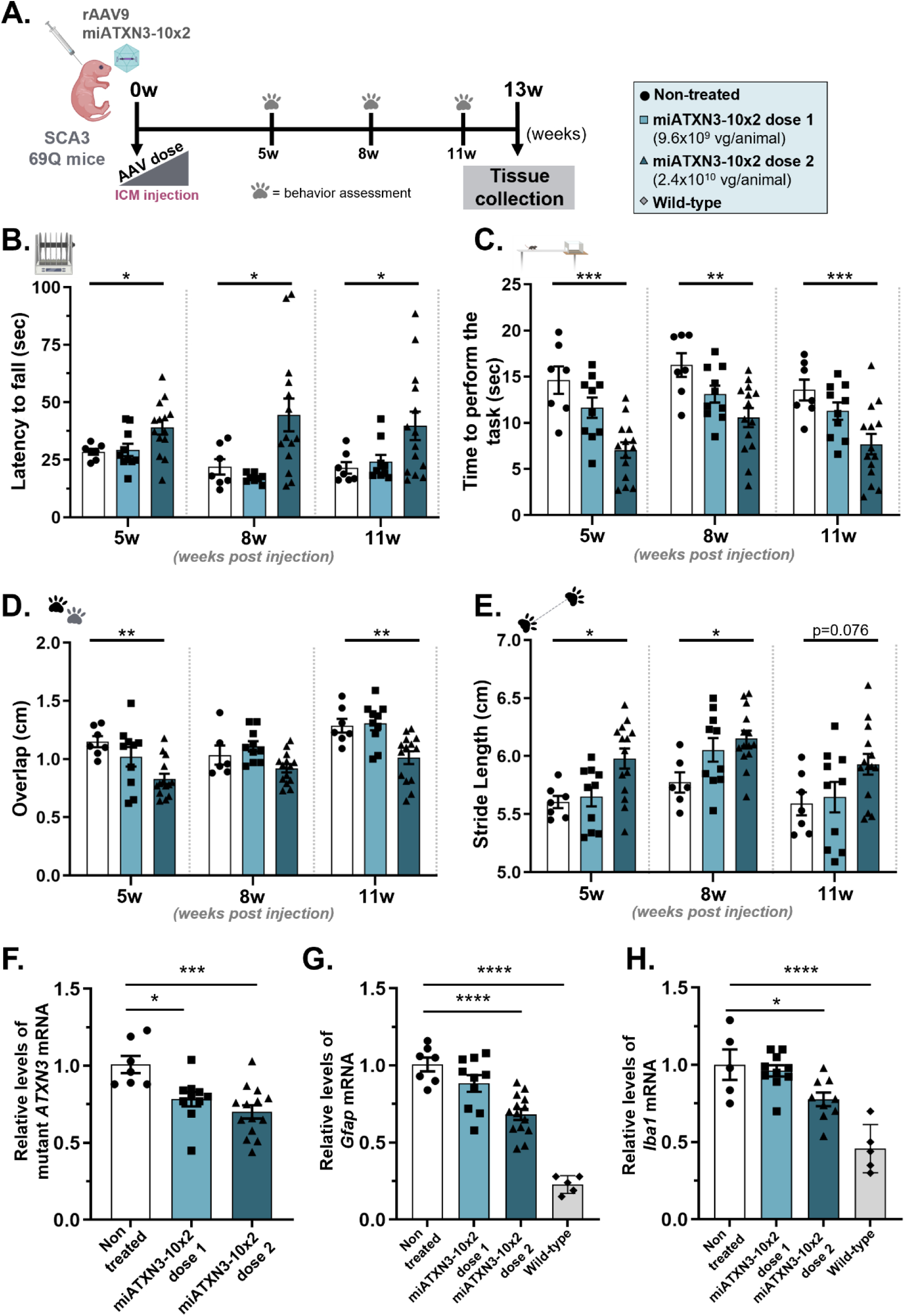
rAAV9-miATXN3-10x2 treatment ameliorates motor phenotype and molecular changes in a severely ataxic SCA3 mouse model following CSF-directed administration. **(A)** Schematic representation of the experimental design and timeline. A long-term efficacy study was conducted using two different doses of rAAV9 vectors packaged with the miATXN3-10x2 silencing construct. Therapeutic vectors were administered via ICM at PND1 in SCA3 transgenic 69Q animals. Motor behavior performance was assessed at three distinct time points – 5, 8 and 11 weeks-of-age – during the study. **(B-E)** Motor coordination, balance, and gait were evaluated, using the rotarod test **(B)**, the beam walking test **(C)** and gait performance test **(D)**. Mice treated with the highest dose of rAAV9-miATXN3-10x2 demonstrated significant improvements in motor performance compared to non-treated animals. Statistical analysis was performed for each time point using the One Way ANOVA with Dunnett’s post-hoc test. **(F-H)** At 13 weeks of age, SCA3 69Q transgenic mice and wild-type littermates were euthanized, and the right brain hemisphere was collected to evaluate the levels of mRNA expression in the cerebellum. A strong and dose-dependent reduction of mutant *ATXN3* mRNA **(F)**, and neuroinflammatory markers, including *Gfap* (G) and Iba1 **(H)** was observed. N=5-14 mice per experimental group. Data presented as mean ±.SEM Statistical analysis was performed using One-way ANOVA, followed by Dunnett’s multiple comparisons post-hoc test, and Pearson’s correlation test (two-tailed p value; grey area denotes the 95% confidence interval).

Motor coordination and balance were first assessed using a rotarod apparatus. While animals treated with the lower dose of miATXN3-10x2 (dose 1) showed no significant improvement in latency to fall compared to non-treated controls, those treated with the highest dose (dose 2) displayed a significant increase at 5, 8, and 11 weeks of age (5 weeks - non-treated: 28.50 sec ± 1.26, miATXN3-10x2 dose 1: 29.26 sec ± 2.67 and miATXN3-10x2 dose 2: 39.07 sec ± 3.12; 8 weeks - non-treated: 21.93 sec ± 3.36, miATXN3-10x2 dose 1: 17.08 sec ± 0.7 and miATXN3-10x2 dose 2: 44.44 sec ± 7.11; 11 weeks - non-treated: 21.49 sec ± 2.49, miATXN3-10x2 dose 1: 24.13 sec ± 2.96 and miATXN3-10x2 dose 2: 39.71 sec ± 6.19) (**Figure 4B**).

To further evaluate motor coordination and balance, the beam-walking test was performed. Similar to the rotarod results, despite no statistically significant differences observed between non-treated and miATXN3-10x2 dose 1 groups, mice treated with the highest dose exhibited significantly lower times to traverse the beam at 5, 8 and 11 weeks of age when compared with control animals (5 weeks - non-treated: 14.61 sec ± 1.48, miATXN3-10x2 dose 1: 11.64 sec ± 1.10 and miATXN3-10x2 dose 2: 7.04 sec ± 0.83; 8 weeks - non-treated: 16.24 sec ± 1.28, miATXN3-10x2 dose 1: 13.10 sec ± 0.91 and miATXN3-10x2 dose 2: 10.56 sec ± 1.03; 11 weeks - non-treated: 13.54 sec ± 1.13, miATXN3-10x2 dose 1: 11.27 sec ± 0.93 and miATXN3-10x2 dose 2: 7.69 sec ± 1.10) (**Figure 4C**).

Finally, gait parameters, including paw overlap and stride length, were assessed using an automated gait analysis system at 5, 8 and 11 weeks of age. A significant decrease in the overlap distance was observed in the miATXN3-10x2 dose 2 group at 5 and 11 weeks when compared to the non-treated controls (5 weeks - non-treated: 1.15 cm ± 0.05, miATXN3- 10x2 dose 1: 1.02 cm ± 0.08 and miATXN3-10x2 dose 2: 0.83 cm ± 0.04; 8 weeks - non-treated: 1.04 cm ± 0.08, miATXN3-10x2 dose 1: 1.11 cm ± 0.04 and miATXN3-10x2 dose 2: 0.92 cm ± 0.04; 11 weeks - non-treated: 1.29 cm ± 0.06, miATXN3-10x2 dose 1: 1.31 cm ± 0.06 and miATXN3-10x2 dose 2: 1.01 cm ± 0.05) (**Figure 4D**).

Stride length, defined as the average distance between consecutive steps for each paw, was also measured. The mean distance between each stride for the hind and front left paws, along with hind and front right paws, was calculated. Impaired SCA3 mice typically exhibited reduced stride lengths compared to wild-type controls. Once again, treatment with dose 2 significantly improved stride length compared with non-treated animals (5 weeks - non-treated: 5.60 cm ± 0.05, miATXN3-10x2 dose 1: 5.65 cm ± 0.08 and miATXN3-10x2 dose 2: 5.98 cm ± 0.09; 8 weeks - non-treated: 5.77 cm ± 0.09, miATXN3-10x2 dose 1: 6.05 cm ± 0.10 and miATXN3-10x2 dose 2: 6.15 cm ± 0.07; 11 weeks - non-treated: 5.59 cm ± 0.10, miATXN3-10x2 dose 1: 5.65 cm ± 0.13 and miATXN3-10x2 dose 2: 5.93 cm ± 0.09) (**Figure 4E**).

Thirteen weeks after vector administration, molecular analysis showed a dose-dependent reduction of mutant *ATXN3* mRNA levels in the cerebellum compared with non-treated controls (non-treated: 1.0 ± 0.06, miATXN3-10x2 dose 1: 0.78 ± 0.05 and miATXN3-10x2 dose 2: 0.70 ± 0.04) (**Figure 4F**). Animals treated with the highest dose presented a reduction of 30% in mutant *ATXN3* transcript levels, which was consistent with the higher levels of vector DNA and miATXN3 copies detected in this group.

Neuroinflammatory markers, including *Gfap* and *Iba1* mRNA levels, which are markedly elevated in the SCA3 69Q animal model, were also analyzed in the cerebellum. Treatment with the highest dose of rAAV9-miATXN3-10x2 significantly reduced *Gfap* mRNA levels and trended towards the levels in wild-type animals (non-treated: 1.0 ± 0.04, miATXN3-10x2 dose 1: 0.88 ± 0.05, miATXN3-10x2 dose 2: 0.68 ± 0.03 and wild-type: 0.23 ± 0.03) (**Figure 4G**). Similar patterns of expression were observed for the Iba1 transcript (non-treated: 1.0 ± 0.09, miATXN3-10x2 dose 1: 0.96 ± 0.04, miATXN3-10x2 dose 2: 0.77 ± 0.04 and wild-type: 0.46 ± 0.07) (**Figure 4H**).

No significant differences in body weight were observed between non-treated and treated animals at any evaluated timepoint (vehicle: 20.91 ± 0.86, miATXN3-10x2 dose 1: 20.61 ± 1.06 and miATXN3-10x2 dose 2: 20.03 ± 0.89) (**Supplementary Figure 3A**). Pearson’s correlation analysis using a correlation matrix of multiple variables from the efficacy study was also performed. Overall, a strong relationship between the number of vector genomes in the cerebellum, mutant *ATXN3* mRNA silencing and motor behavior performance was detected (**Supplementary Figure 3B**). This suggests that robust dose-response outcomes were observed following treatment with our therapeutic strategy.

Moreover, no significant differences in rotarod performance were observed in wild-type animals treated with dose 1 and dose 2 of rAAV9-miATXN3-10x2 at 11 weeks of age (**Supplementary Figure 4**), suggesting no overt neurotoxic effects.

Collectively, these findings highlight the therapeutic potential of rAAV9-miATXN3-10x2 administered via ICM injection, demonstrating significant improvements in motor performance, target engagement, and partial correction of the dysregulated glial markers in a severely impaired SCA3 mouse model.

### rAAV9-miATXN3-10x2 reduces mutant ATXN3 aggregation and mitigates cerebellar degeneration in the SCA3 69Q mouse model

A hallmark pathological feature of SCA3 is the formation of neuronal inclusions composed of aggregated mutant ATXN3 protein (Paulson et al., 1997). In accordance, the SCA3 69Q mouse model develops mutant ATXN3 protein aggregates in the cerebellum. To evaluate whether rAAV9-miATXN3-10x2 could reduce aggregation, we performed immunolabeling for the HA tag fused to the mutant ATXN3 protein in the SCA3 69Q animal model (**Figure 5A**). The number of mutant ATXN3 aggregates per mm^2^ was markedly lower in cerebellar lobules of animals treated with rAAV9-miATXN3-10x2 than in non-treated controls (non-treated: 89.60 ± 14.57, miATXN3-10x2 dose 1: 41.75 ± 2.46 and miATXN3-10x2 dose 2: 26.89 ± 1.81) (**Figure 5B**).

**Figure 5.**
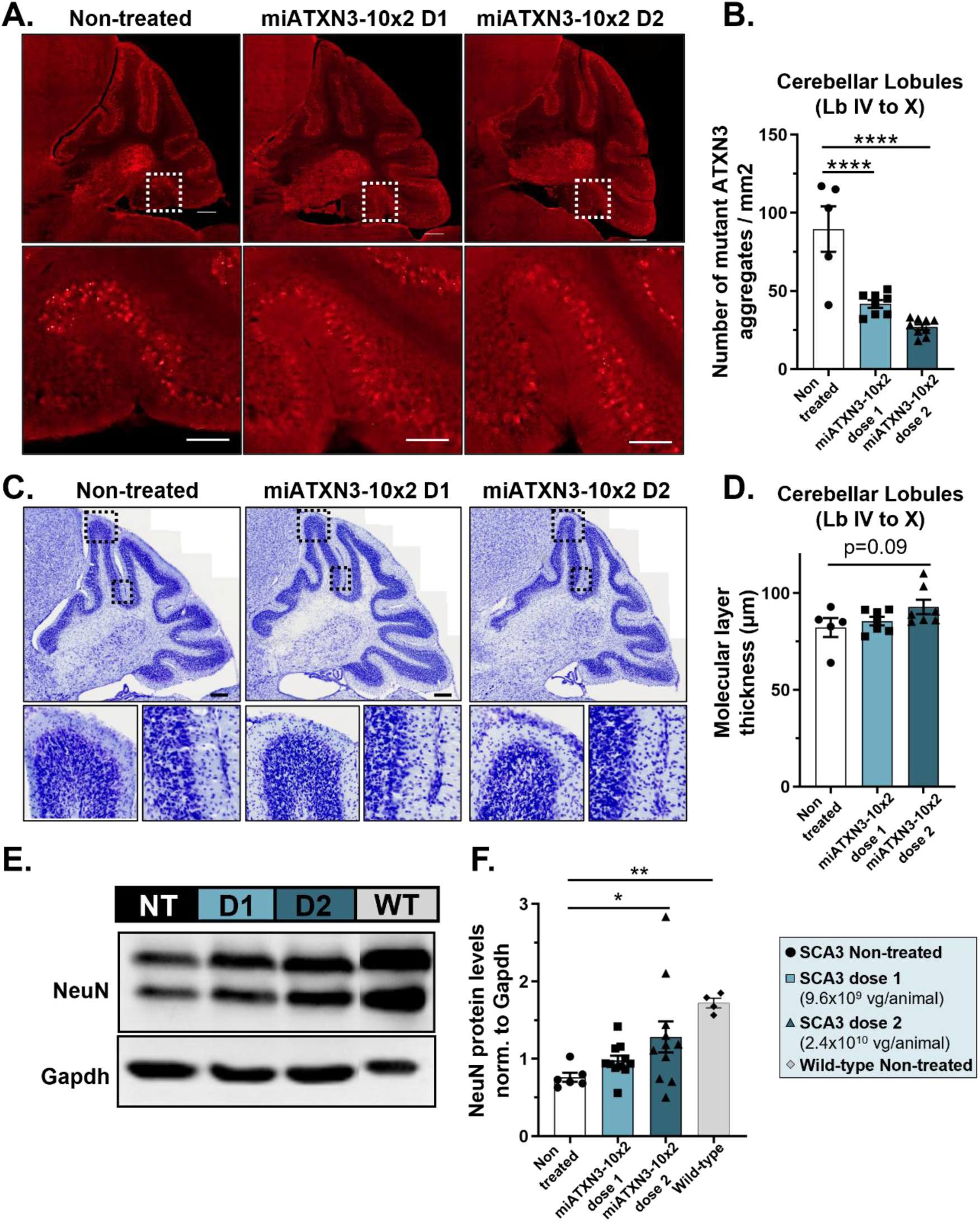
rAAV9-miATXN3-10x2 administration via ICM injection alleviates neuropathological alterations in the severely impaired transgenic SCA3 69Q mouse model. **(A)** Representative immunofluorescent images showing HA-tagged mutant ATXN3 aggregates in the cerebellum of 13-week-old vehicle- or rAAV9-miATXN3-10x2-treated animals. **(B)** Quantification of mutant ATXN3 aggregates in cerebellar lobules IV to X. Treated animals showed a significant reduction in the number of mutant ATXN3 aggregates compared with controls. **(C)** Representative images of cresyl violet staining in the cerebellum of 13 weeks-old vehicle- or rAAV9-miATXN3- 10x2-treated animals. **(D)** Quantification of molecular layer thickness in cerebellar lobules IV to X. **(E)** Western blot analysis of the neuronal marker NeuN in the cerebellum of treated mice. **(F)** Restoration of NeuN protein levels was observed in animals that received the highest dose of rAAV9-miATXN3-10x2. N=4-14 mice per experimental group. Data presented as mean± SEM. Statistical analysis was performed using One-way ANOVA, followed by Dunnett’s multiple comparisons post-hoc test. Outliers were removed according to the Grubbs test (α=0.05). NT - non-treated SCA3 69Q transgenic, D1 - SCA3 69Q transgenic receiving dose 1 of rAAV9-miATXN3-10x2, D2 - SCA3 69Q transgenic receiving dose 2 of rAAV9-miATXN3-10x2, WT – non-treated wild-type.

Additionally, shrinkage of the cerebellum is a characteristic feature of pathology in the SCA3 brain, including gray matter atrophy (Goel et al., 2011; Scherzed et al., 2012; Peng et al., 2019; Guo et al., 2020; Miranda et al., 2022). Therefore, to investigate the potential neuroprotective effects of rAAV9-miATXN3-10x2, we first measured the thickness of the molecular layer in the cerebellum. Animals treated with the highest dose of rAAV9-miATXN3-10x2 showed a trend toward increased molecular layer thickness, suggesting a protective effect against cerebellar layer shrinkage (non-treated: 82.24 ± 4.86, miATXN3- 10x2 dose 1: 85.53 ± 2.17 and miATXN3-10x2 dose 2: 92.88 ± 3.73) (**Figure 5C and D**). To determine whether rAAV9-miATXN3-10x2 could also mitigate neuronal loss, we measured NeuN protein levels - a marker of neuronal integrity - in cerebellar lysates by western blot analysis (**Figure 5E**). Treated animals displayed a dose-dependent increase in NeuN levels compared to non-treated controls. This increase was statistically significant in animals receiving the highest dose, with levels approaching those observed in wild-type animals (non-treated: 0.76 ± 0.06, miATXN3-10x2 dose 1: 0.98 ± 0.06, miATXN3-10x2 dose 2: 1.28 ± 0.20 and wild-type: 1.72 ± 0.06) (**Figure 5F**).

Together, these results demonstrate that rAAV9-miATXN3-10x2 effectively prevents mutant ATXN3 aggregation, preserves cerebellar molecular layer thickness, and reduces neuronal cell loss, highlighting its therapeutic potential in addressing neuropathological deficits associated with SCA3.

### rAAV9-miATXN3-10x2 treatment promotes beneficial transcriptomic changes in the cerebellum of SCA3 69Q mice

To evaluate whether our therapeutic strategy could improve the transcriptomic profile of SCA3 69Q mice, we performed RNA sequencing on cerebellar samples collected at 13 weeks of age following ICM delivery of rAAV9-miATXN3-10x2. Samples were obtained from non-treated SCA3 69Q animals, treated SCA3 69Q animals that achieved >30% silencing of mutant *ATXN3* mRNA (referred to as the “top treated group”), and non-treated wild-type littermates (**Figure 6A**). Differentially expressed genes (DEGs) were identified by comparing non-treated transgenic animals with the top-treated group. A volcano plot of these comparisons revealed 2009 DEGs, comprising 1133 upregulated and 876 downregulated genes in top-treated animals relative to non-treated controls, demonstrating that rAAV9-treatment induced significant alterations in gene expression (**Figure 6B**).

**Figure 6.**
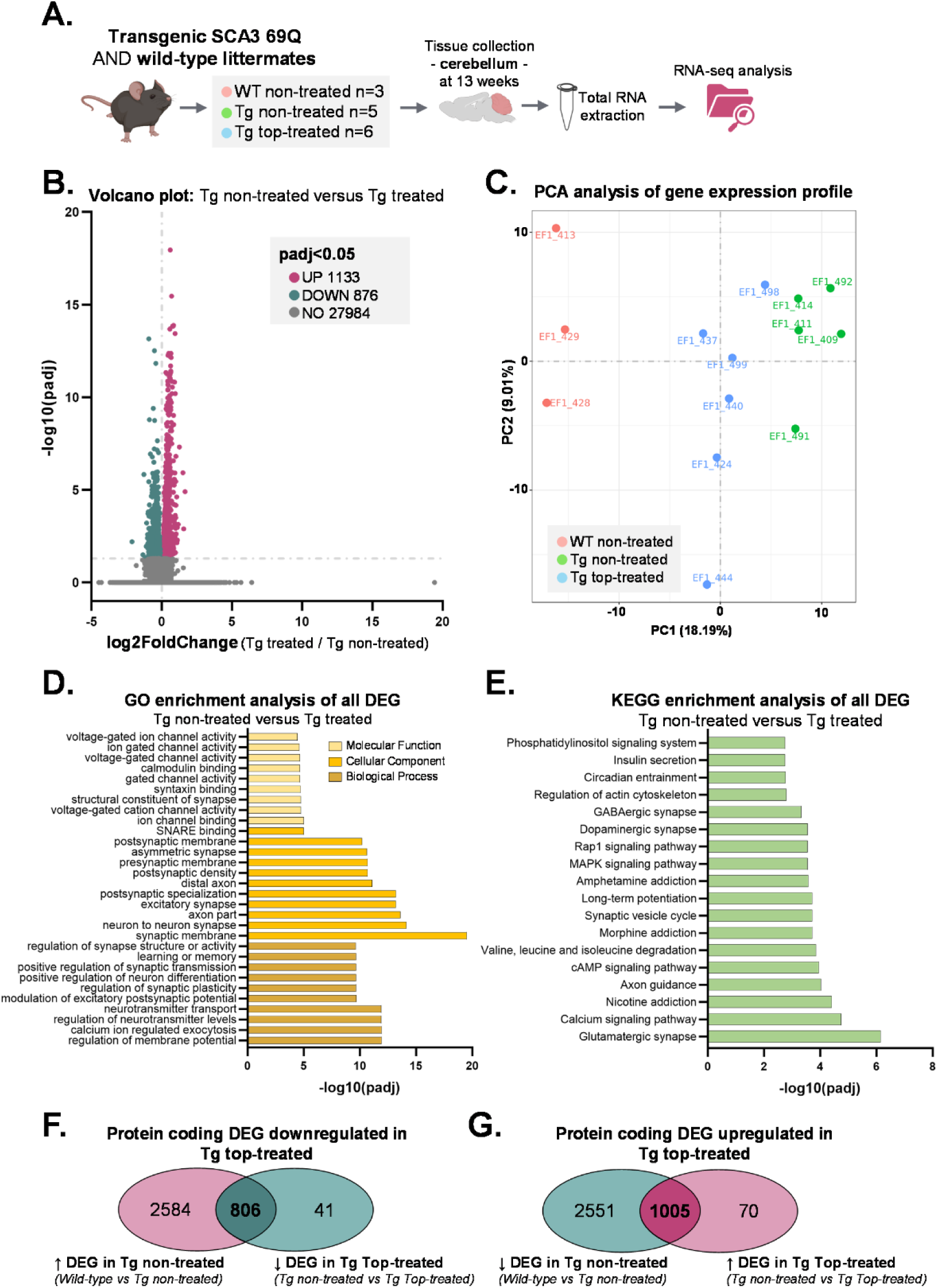
Administration of rAAV9-miATXN3-10x2 improves the cerebellar transcriptomic profile of a the SCA3 69Q mouse model. **(A)** Schematic representation of the experimental design. Transgenic (Tg) SCA3 69Q mice from the efficacy study, either non-treated or top treated with rAAV9-miATXN3-10x2 (>30% silencing of human mutant ATXN3 mRNA), along with wild-type (WT) littermates, were selected for transcriptome analysis of cerebellar tissue at 13 weeks of age. **(B)** Volcano plot displaying differentially expressed genes (DEGs) in non-treated transgenic versus top-treated transgenic animals. A total of 2009 genes showed significant expression changes in top-treated animals compared to non-treated controls, with 1133 genes being upregulated (UP) and 876 downregulated (DOWN). **(C)** Principal component analysis (PCA) analysis of mRNA sequencing data from non-treated wild-type, non-treated transgenic and top-treated transgenic animals. **(D)** Gene Ontology (GO) enrichment analysis highlighting the most overrepresented GO terms among all DEGs, including upregulated and downregulated genes. **(E)** Kyoto Encyclopedia of Genes and Genomes (KEGG) pathway analysis identifying the top enriched pathways among all DEGs. **(F-G)** Venn diagrams analysis showing the overlap of protein coding DEGs between non-treated transgenic animals versus wild-type littermates, as well as between non-treated and top-treated transgenic animals. Potentially rescued genes, both downregulated **(F)** and upregulated **(G)** in top-treated animals are highlighted.

To determine whether these transcriptomic changes were beneficial, we performed Principal Component Analysis (PCA) of the mRNA sequencing data. The PCA revealed clustering of treated animals closer to the wild-type cluster, suggesting that rAAV9-miATXN3-10x2 treatment shifted the transcriptomic profile of SCA3 69Q mice toward a wild-type-like state (**Figure 6C**).

We further explored the biological significance of these changes using Gene Ontology (GO) enrichment analysis of all DEGs, including both upregulated and downregulated genes in the top-treated group (**Figure 6D**). The most enriched GO terms were related to the synaptic membrane, neuron-to-neuron synapse, axon and excitatory synapse. Other overrepresented terms were associated with the modulation of synaptic transmission, including synaptic structure, function, and channel activity. Kyoto Encyclopedia of Genes and Genomes (KEGG) analysis further highlighted the enrichment of pathways related to synaptic function and neuronal signaling. These included glutamatergic, dopaminergic, and GABAergic synapse pathways, as well as signaling pathways critical to neuronal health, such as calcium signaling, cAMP signaling, MAPK signaling and Rap1 signaling pathways (**Figure 6E**).

Finally, we investigated whether the genes that were differentially expressed in top-treated animals could potentially be rescued to levels comparable to wild-type mice. Among protein-coding DEGs, we identified 3390 genes that were upregulated in non-treated transgenic animals relative to wild-type controls and found that 806 of these genes were downregulated in top-treated transgenic animals compared to non-treated transgenic animals, reflecting a shift toward wild-type expression levels (**Figure 6F**). Similarly, among the 3556 downregulated protein-coding DEGs in non-treated transgenic animals relative to wild-type controls, 1005 of these genes were upregulated in top-treated transgenic animals compared to non-treated transgenic animals (**Figure 6G**). Altogether, these results indicate that a total of 1811 protein-coding genes exhibited expression changes trending toward wild-type levels following rAAV9-miATXN3-10x2 treatment.

In summary, these findings demonstrate that rAAV9-miATXN3-10x2 administration induces significant and beneficial transcriptomic changes in the cerebellum of SCA3 69Q mice, including the rescue of key neuronal pathways and the restoration of gene expression profiles closer to those of wild-type animals.

## DISCUSSION

SCA3 is a severe and fatal neurodegenerative disorder for which no effective treatment currently exists to alter its progression. The existing options rely on symptom management through pharmacotherapy and physiotherapy, leaving a significant unmet medical need for patients and caregivers. Given its monogenic nature, SCA3 is an ideal candidate for molecular therapies aiming at silencing the mutant *ATXN3* mRNA, thereby targeting the disease at the root cause of toxicity and preventing downstream neurodegenerative processes.

RNAi, combined with viral vector delivery, has emerged as a powerful tool for sustained gene silencing in preclinical and clinical settings. In this study, we optimized a therapeutic strategy using rAAVs encoding artificial miRNAs designed to specifically target the mutant *ATXN3* mRNA. Allele-specific silencing was achieved by targeting intragenic SNPs in linkage disequilibrium with the expanded *ATXN3* allele. Specifically, our approach focused on SNP8 (rs1048755) and SNP10 (rs12895357) variants associated with the mutant allele in 72% of SCA3 patients (Gaspar et al., 2001). While alternative strategies, such as targeting of the expanded CAG repeats, are being explored in clinical trials (reviewed in de Sousa-Lourenço et al., 2024), they carry the risk of off-target effects due to the presence of CAG repeats in multiple genes throughout the human genome (Butland et al., 2007). In contrast, allele-specific silencing via SNPs minimizes such risks and preserves the function of wild-type ATXN3 protein, which plays critical roles in cellular processes, including deubiquitination, transcriptional regulation, and DNA repair (reviewed in Matos et al., 2019). Indeed, our group and others have achieved mutant *ATXN3* allele-specific silencing via SNP targeting in different SCA3 models (Alves et al., 2008; Nóbrega et al., 2013, 2014; Conceição et al., 2016; Nobre & Pereira de Almeida, 2020; Hauser et al., 2022; Rufino-Ramos et al., 2023).

Our *in vitro* experiments demonstrated that the bicistronic miRNA construct miATXN3-10x2, targeting SNP10 (rs12895357) achieved potent and specific silencing of the mutant *ATXN3* mRNA, while maintaining normal levels of wild-type *ATXN3* mRNA. The use of bicistronic miRNAs has been hypothesized to enhance silencing potency, potentially enabling lower vector doses to achieve therapeutic efficacy. This is a critical consideration, particularly in light of the dose-dependent toxicity observed in non-human primates at high rAAV doses (Hinderer et al., 2018; Hordeaux et al., 2020).

The lead candidate miATXN3-10x2 was then packaged into rAAV9 vectors for *in vivo* target engagement studies. Our experiments in two distinct SCA3 mouse models (YAC84Q and 69Q) confirmed the therapeutic potential of rAAV9-miATXN3-10x2. Intra-cerebellar administration in both animal models resulted in efficient transduction, as well as significant reduction of mutant ATXN3 mRNA and protein levels in the cerebellum. However, while mutant *ATXN3* mRNA was suppressed in the brainstem, protein levels remained unchanged, likely due to insufficient vector distribution in this brain region. These findings highlight the challenges associated with localized parenchymal delivery and underscore the need for alternative administration routes that can achieve broader CNS targeting.

To address this, we explored ICM injection as a less invasive delivery method for broad CNS biodistribution and clinical translatability. First, ICM administration of rAAV9 encoding the GFP reporter in neonatal SCA3 YAC84Q mice resulted in widespread GFP expression throughout the brain, including the cerebellum and brainstem. Our findings aligned with previous studies that reported rAAV9 efficient transduction throughout the brain and spinal cord following a single ICM administration in neonatal mice (Lukashchuk et al., 2016; Wiseman et al., 2024) and adult non-human primates (Hinderer et al., 2014; Matsuzaki et al., 2017). Moreover, rAAV9 injection into the cisterna magna of cynomolgus macaques was shown to be 100- and 10-fold more efficient at transducing the brain and spinal cord, respectively, compared to lumbar intrathecal administration (Hinderer et al., 2014). This highlights the advantages of ICM injection for simultaneously targeting both the spinal cord and deeper brain structures. This approach also circumvents issues related to circulating neutralizing antibodies and hepatotoxicity associated with systemic rAAV delivery (Gray et al., 2013; Hinderer et al., 2018; reviewed in Daci & Flotte, 2024). Although ICM administration poses technical challenges in humans, advancements such as fluoroscopic-guided lumbar access to the cisterna magna (Taghian et al., 2020) offer promising solutions for clinical translation.

In a long-term efficacy study, ICM delivery of rAAV9-miATXN3-10x2 in SCA3 69Q mice led to sustained improvement in motor coordination, balance, and gait parameters. In fact, mice treated with the highest dose achieved approximately a 2-fold increase in rotarod performance and a 2-fold reduction in the time to cross the beam.

A key hallmark of SCA3 is the accumulation of mutant ATXN3 inclusions in the affected brain regions (Paulson et al., 1997). Furthermore, significant neuronal loss, including in the Purkinje cell layer and DCNs (Scherzed et al., 2012), along with a reduction in cerebellar volume (Peng et al., 2019; Wan et al., 2020; Miranda et al., 2022), have been observed in the SCA3 brain. These neuropathological changes are accompanied by the upregulation of inflammatory genes and activation of glial cells (Evert et al., 2001; Rüb et al., 2002; Rub et al., 2002; Shi et al., 2015; Switonski et al., 2015).

Remarkably, in the long-term efficacy study, the observed behavioral improvements were accompanied by significant reductions in mutant *ATXN3* mRNA levels (up to 30%) and neuroinflammatory markers (Iba1 and *Gfap*), along with a decrease in mutant ATXN3 protein aggregates and reduced neuronal loss. Importantly, no overt neurotoxicity was observed in treated wild-type animals, further supporting the safety of this approach.

Transcriptomic analysis revealed that rAAV9-miATXN3-10x2 treatment partially rescued the cerebellar gene expression profile in SCA3 69Q mice, shifting it toward a wild-type-like state. Enrichment of synaptic and neuronal signaling pathways, along with the partial restoration of over 1800 protein-coding genes, suggests that this therapy not only reduces mutant *ATXN3* expression but also restores key neuronal functions. These findings provide mechanistic insights into the therapeutic benefits observed at the molecular, cellular, and behavioral levels.

In summary, our study demonstrates the efficacy of rAAV9-miATXN3-10x2 as a potent and allele-specific gene silencing therapy for SCA3. Both intra-cerebellar and ICM delivery methods resulted in robust suppression of mutant *ATXN3* mRNA, amelioration of neuropathological deficits, and significant improvements in motor function. In particular, ICM administration offers a clinically feasible route for broad CNS targeting, paving the way for translating this therapeutic strategy to SCA3 patients. These findings also highlight the broader potential of rAAV-based gene silencing therapies for monogenic neurodegenerative disorders.

## AUTHOR CONTRIBUTIONS

L.P.A., R.J.N., A.C.S, C.H., A.M., S.d.M., M.W., J.N., A.B., and D.L. conceived and designed experiments. A.C.S, C.H., D.L., A.R.F., M.M.L., K.L, D.P. and S.P.D. performed the experiments. A.C.S, C.H., D.L., A.R.F., M.M.L., K.L., S.M.L., M.S., A.M., S.d.M., M.W., J.N., A.B., R.J.N. and L.P.A performed data analysis. A.C.S. and R.J.N. wrote the first draft of the manuscript. All authors reviewed and edited the final manuscript.

## FUNDING AND ACKNOWLEDGMENTS

This work was partially funded by PTC Therapeutics. Our group is also supported by the European Regional Development Fund (ERDF) through the Centro 2020 Regional Operational Programme, the Operational Programme for Competitiveness and Internationalisation (COMPETE 2020), and by Portuguese national funds via Fundação para a Ciência e a Tecnologia (FCT), under the projects UIDB/04539/2025, UIDP/04539/2025, LA/P/0058/2020 and 2022.06118.PTDC; as well as by the projects SpreadSilencing (POCI-01-0145-FEDER029716), ViraVector (CENTRO-01-0145-FEDER-022095) and Neurodiet (JPND/0001/2022). Additional support was provided by CinTech under PRR, ARDAT under the IMI2 JU Grant agreement No 945473 (co-funded by EU and EFPIA), and Capacity 2023 (ID: 101145599), GeneT (ID: 101059981), ERDERA (ID: 101156595), GeneH (ID: 101186939) and GCure (ID: 101186929), under the European Union’s Horizon Europe program. Further funding was received from the American Portuguese Biomedical Research Fund (APBRF), the European Advanced Translational Research Infrastructure for Neurosciences (NeurATRIS), and the Richard Chin and Lily Lock MJD Research Fund. FCT PhD fellowships supported the work of A.C.S. (2020.07721.BD), C.H. (2021.06939.BD), D.L. (2020.09668.BD), A.R.F. (2021.08215.BD), M.M.L. (2021.05776.BD) and K.L. (2020.09513.BD). We also thank all members of the L.P.A. laboratory for the support and discussions.

**See Supplemental material for supplemental figures and tables.**

