## Supplemental material for "rAAV-Delivered Bicistronic Artificial microRNAs for Allele-Specific Silencing Improve Motor and Molecular Outcomes in Spinocerebellar Ataxia Type 3"

**Author names and affiliations:**

Ana Carolina Silva[1-5,*], Carina Henriques[1,2,4-6,*], Diana D. Lobo[1-5,*], Ana Rita Fernandes[1-4], Miguel M. Lopes[1-5], Kevin Leandro[1,2,4,6], Dina Pereira[1-4], Sónia P. Duarte[1-4], Sara M. Lopes[1-4], Magda M. Santana[1-4], Amal Dakka[7], Steve De Marco[7], Marla Weetall[7], Jana Narasimhan[7], Anu Bhattacharyya[7], Rui Jorge Nobre[1-5,+], Luís Pereira de Almeida[1,2,4-6,+]

1. Center for Neuroscience and Cell Biology (CNC), Gene and Stem Cell Therapies for the Brain Group, University of Coimbra, 3004-504 Coimbra, Portugal

2. Center for Innovative Biomedicine and Biotechnology (CIBB), Vectors, Gene and Cell Therapy Group, University of Coimbra, 3004-504 Coimbra, Portugal

3. Institute for Interdisciplinary Research (III), University of Coimbra, 3030-789 Coimbra, Portugal

4. GeneT, Center for Excellence in Gene Therapy in Portugal, University of Coimbra, 3004-504 Coimbra, Portugal

5. ViraVector–Viral Vector for Gene Transfer Core Facility, University of Coimbra, 3004-504 Coimbra, Portugal

6. Faculty of Pharmacy, University of Coimbra, 3000-548 Coimbra, Portugal

7. PTC Therapeutics, Inc. NJ Center of Excellence Campus 1041 U.S. Route 202. Building 1013. Bridgewater, NJ 08807, USA

*Equal contribution as first authors; +Equal contribution as senior authors

**Corresponding Authors:**

Luís Pereira de Almeida and Rui Jorge Nobre

CNC-UC - Center for Neuroscience and Cell Biology, University of Coimbra, Rua Larga, 3004-504 Coimbra, Portugal

### SUPPLEMENTAL MATERIAL:

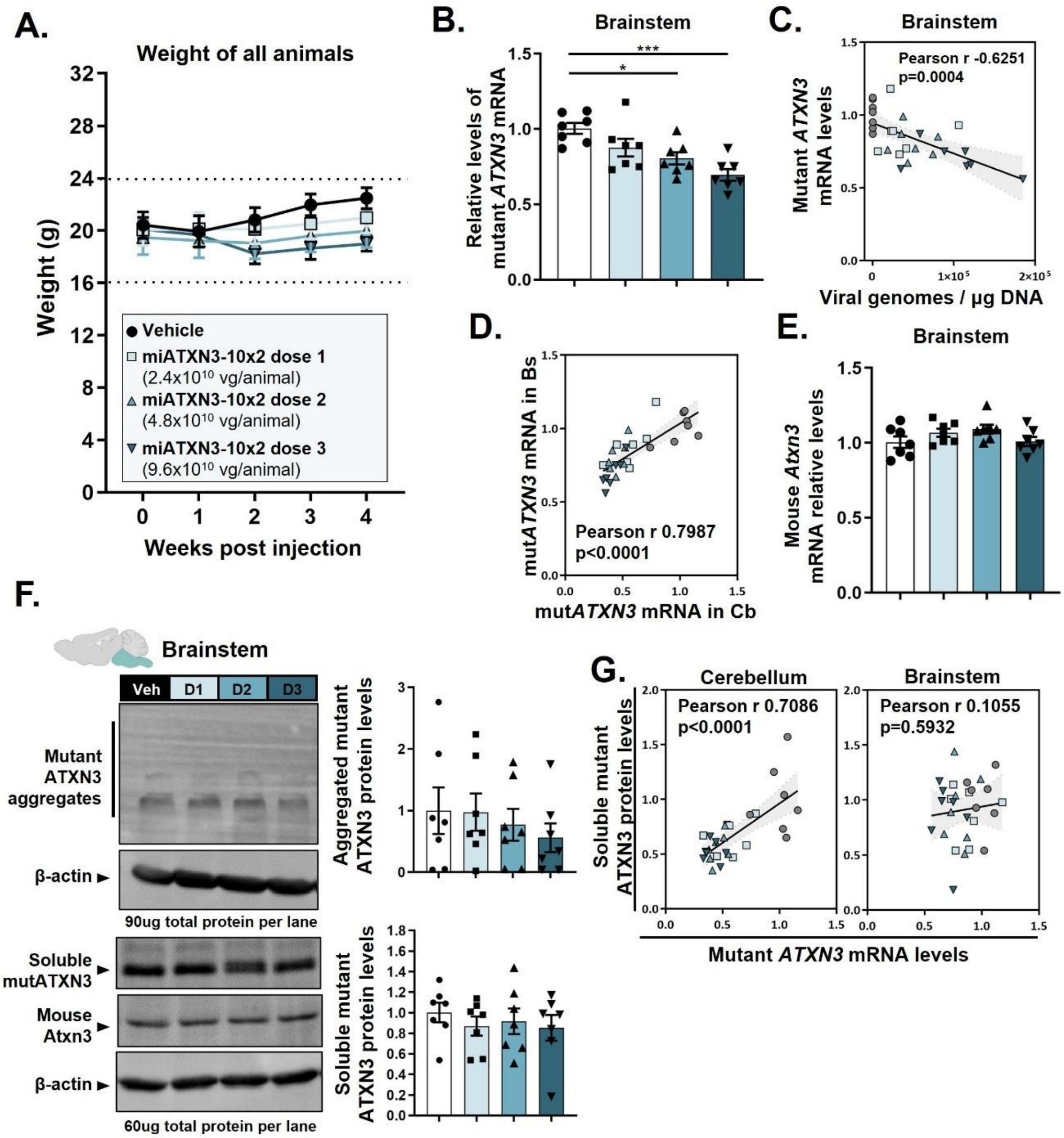


**Supplementary Figure 1. Target engagement study in hemizygous SCA3 YAC84Q mice following rAAV9-miATXN3-10x2 intraparenchymal administration into the cerebellum.** At 10 weeks of age, hemizygous SCA3 YAC84Q mice received bilateral injections in each deep cerebellar nuclei (DCN) with vehicle or rAAV9-miATXN3-10x2 at three different vector doses. **(A)** Body weight was measured weekly following vector administration. Statistical analysis, performed using two-way ANOVA for simple effects followed by Tukey's multiple comparisons test, revealed no significant differences in body weight between treatment groups at any time point evaluated. **(B)** Four weeks following surgery, the mice were euthanized, and brain tissues were collected for analysis. RT-qPCR analysis of the brainstem tissue showed a strong and dose-dependent reduction of mutant ATXN3 mRNA levels. **(C)** A significant inverse correlation between the number of viral genomes and mutant ATXN3 mRNA levels was found in the brainstem. **(D)** Correlation analysis to evaluate the relationship between mutant ATXN3 mRNA levels in the cerebellum and brainstem revealed a strong positive correlation for these parameters. **(E)** No differences in endogenous mouse Atxn3 mRNA levels were observed in the brainstem region. **(F)** Western blot analysis revealed no significant differences in mutant ATXN3 aggregated and soluble protein levels in the brainstem of treated mice. **(G)** Correlation analyses were performed to evaluate the relationship between mutant ATXN3 mRNA levels and soluble mutant ATXN3 protein levels in disease-relevant brain regions, which was found to be statistically significant in the cerebellar tissue. N=7 mice per experimental group. Data presented as mean ± SEM. Statistical analysis was performed using One-way ANOVA, followed by Dunnett's multiple comparisons post-hoc test, and Pearson’s correlation test (two-tailed p value; grey area denotes the 95% confidence interval). Cb - Cerebellum, Bs - Brainstem.

54

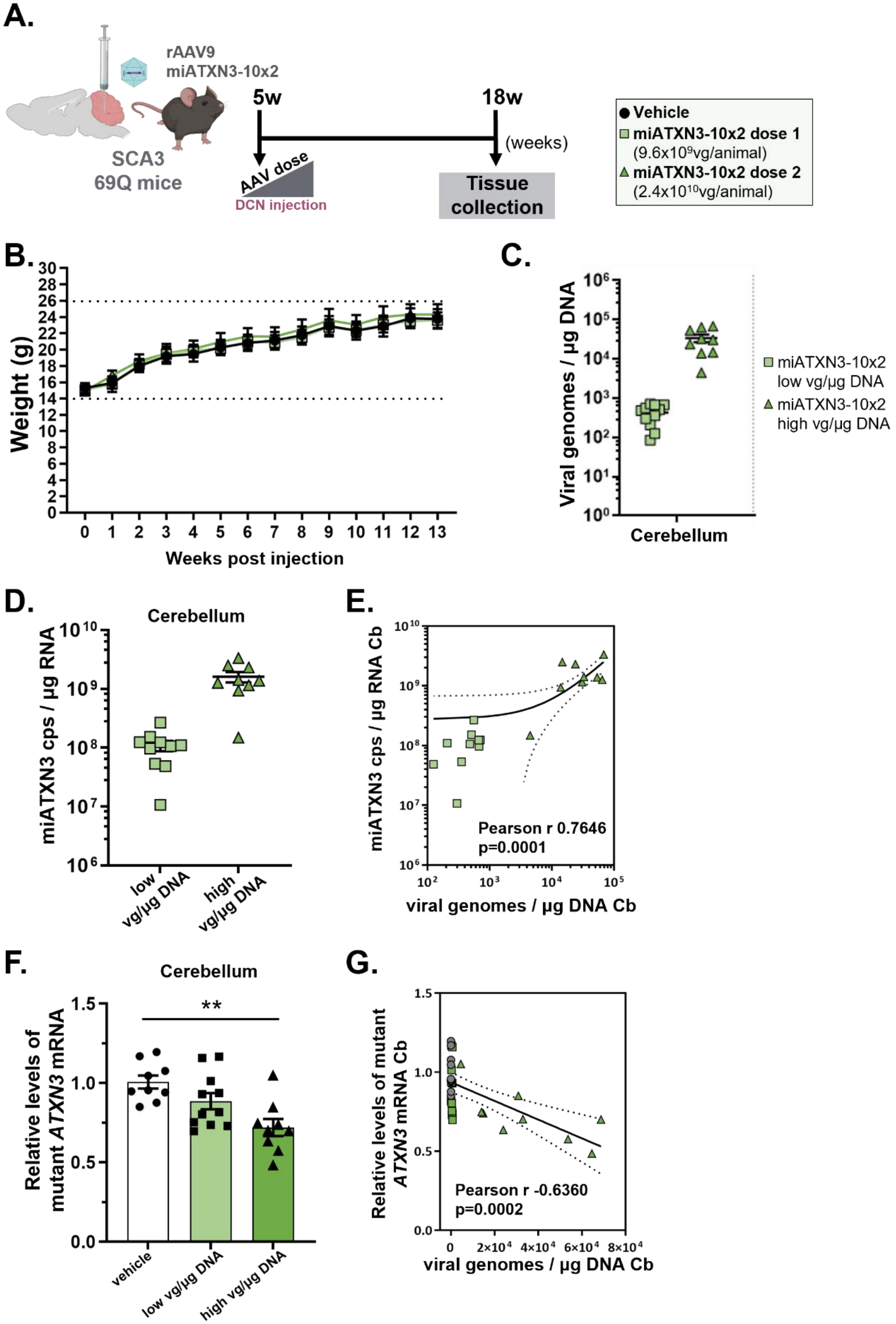
A.
rAAV9
miATXN3-10x2
SCA3
69Q mice
5w
18w
(weeks)
AAV dose
DCN injection
Tissue collection
Vehicle
miATXN3-10x2 dose 1
(9.6x10⁹vg/animal)
miATXN3-10x2 dose 2
(2.4x10¹⁰vg/animal)
B.
Weight (g)
Weeks post injection
C.
Viral genomes / µg DNA
Cerebellum
miATXN3-10x2 low vg/µg DNA
miATXN3-10x2 high vg/µg DNA
D.
Cerebellum
miATXN3 cps / µg RNA
low vg/µg DNA
high vg/µg DNA
E.
miATXN3 cps / µg RNA Cb
Pearson r 0.7646
p=0.0001
viral genomes / µg DNA Cb
F.
Cerebellum
Relative levels of mutant ATXN3 mRNA
**
vehicle
low vg/µg DNA
high vg/µg DNA
G.
Relative levels of mutant ATXN3 mRNA Cb
Pearson r -0.6360
p=0.0002
viral genomes / µg DNA Cb

**Supplementary Figure 2. Target engagement study in transgenic SCA3 69Q mice following direct intra-cerebellar administration of rAAV9-miATXN3-10x2. (A)** Schematic representation of the experimental design and timeline. At 5 weeks of age, SCA3 transgenic 69Q mice expressing a truncated form of expanded human ATXN3 in the cerebellum received bilateral injections into the deep cerebellar nuclei (DCN) with either vehicle or rAAV9-miATXN3-10x2 at two different vector doses. Thirteen weeks post-administration, the mice were euthanized for collection of brains. Total DNA and RNA were extracted and used to perform qPCR analysis. **(B)** Weight of all animals was measured weekly throughout the experimental period. Statistical analysis was performed to assess weight differences between groups at each time-point using the Two-way ANOVA simple effects, followed by Tukey's multiple comparisons test at each time point. No significant body weight differences between groups were observed at each time point evaluated. (**C)** Vector distribution was evaluated by quantification of viral genome copies in the cerebellum. Animals were distributed in two dose groups according to the number of viral genomes that reached the cerebellum (low vg/µg DNA and high vg/µg DNA groups). **(D)** The number of miRNA copies was determined in cerebellar tissue, revealing detectable levels consistent with vector dose that reached this brain region. **(E)** A strong positive correlation was observed between viral genomes and miATXN3 copy numbers in the cerebellum. **(F)** Target engagement was assessed by measuring mutant ATXN3 mRNA levels, demonstrating a significant and dose-dependent reduction in treated animals. **(G)** A strong negative correlation was observed between the number of viral genomes and levels of mutant ATXN3 mRNA in the cerebellum. N=9-11 mice per experimental group. Data presented as mean ± SEM. Statistical analysis was conducted using One-way ANOVA, followed by Dunnett's multiple comparisons post-hoc test, and Pearson's correlation test (two-tailed p value; dotted lines denote the 95% confidence interval). Cb - Cerebellum,

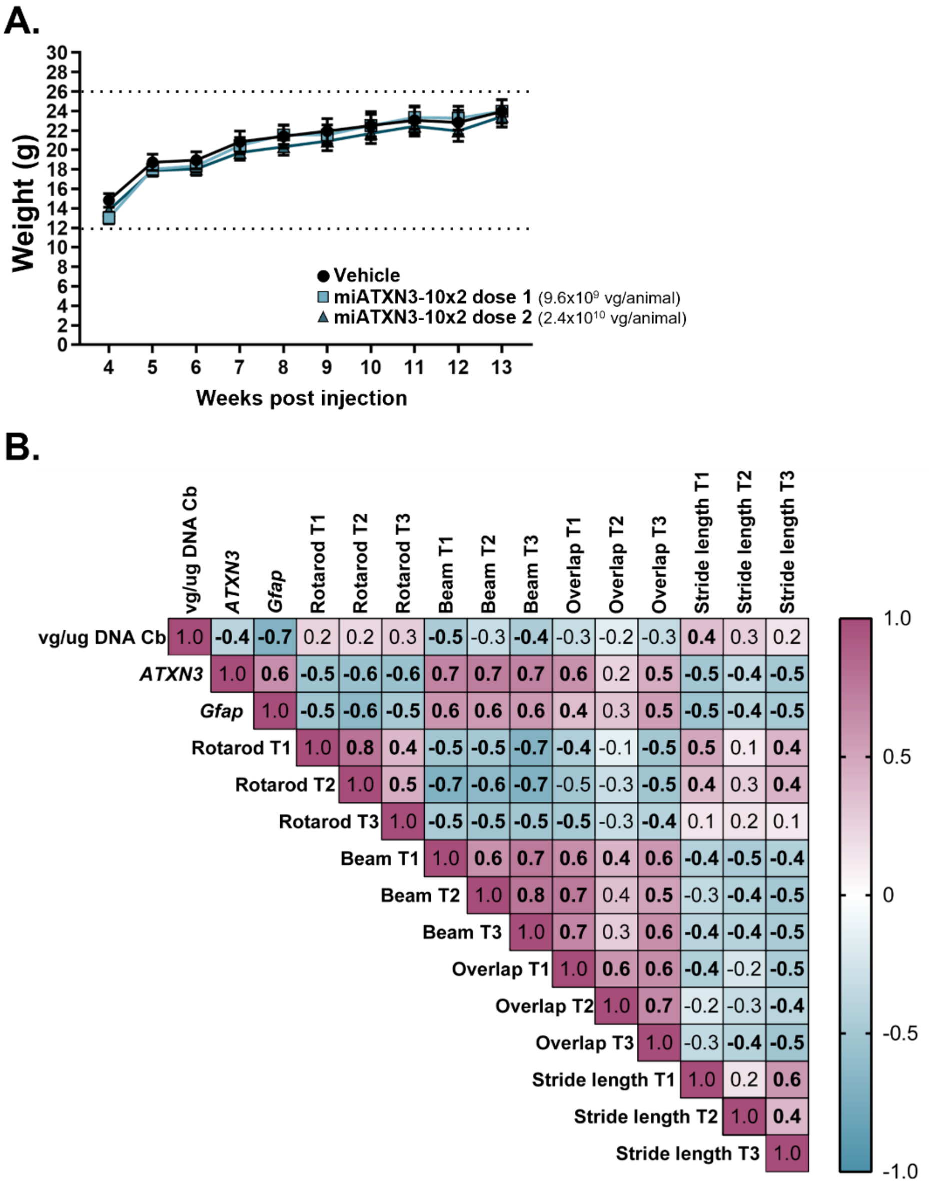


**Supplementary Figure 3. Correlation matrix of multiple variables in the long-term efficacy study following CSF administration of rAAV9-miATXN3-10x2. (A)** Animal weights were recorded weekly following vector administration. No significant differences in body weight were observed between experimental groups at any time point. Statistical analysis was performed using two-way ANOVA with simple effects and Tukey's multiple comparisons test. N=5-14 mice per experimental group. Data presented as mean±sem. **(B)** Pearson's correlation analysis was performed using a correlation matrix of multiple variables from the study. The colored squares show the Pearson's r correlation coefficient between two variables, with statistically significant correlations ($p<0.05$) indicated in bold. Pink shades denote positive correlations, while blue shades show negative correlations, with deeper color representing a stronger correlation.. Vg/ug DNA Cb - number of viral genomes per ug of total DNA in the cerebellum; ATXN3 - Relative levels of mutant ATXN3 mRNA in the cerebellum; Gfap - Relative levels of mutant Gfap mRNA in the cerebellum; Rotarod T1, T2 and T2 - Rotarod test performance in timepoint 1, 2 and 3, respectively; Beam T1, T2 and T2 - Beam walking test in timepoint 1, 2 and 3, respectively. Overlap T1, T2 and T2 - Paw overlap values from gait test performance in timepoint 1, 2 and 3, respectively. Stride length T1, T2 and T2 - Stride length values from gait test performance in timepoint 1, 2 and 3, respectively.

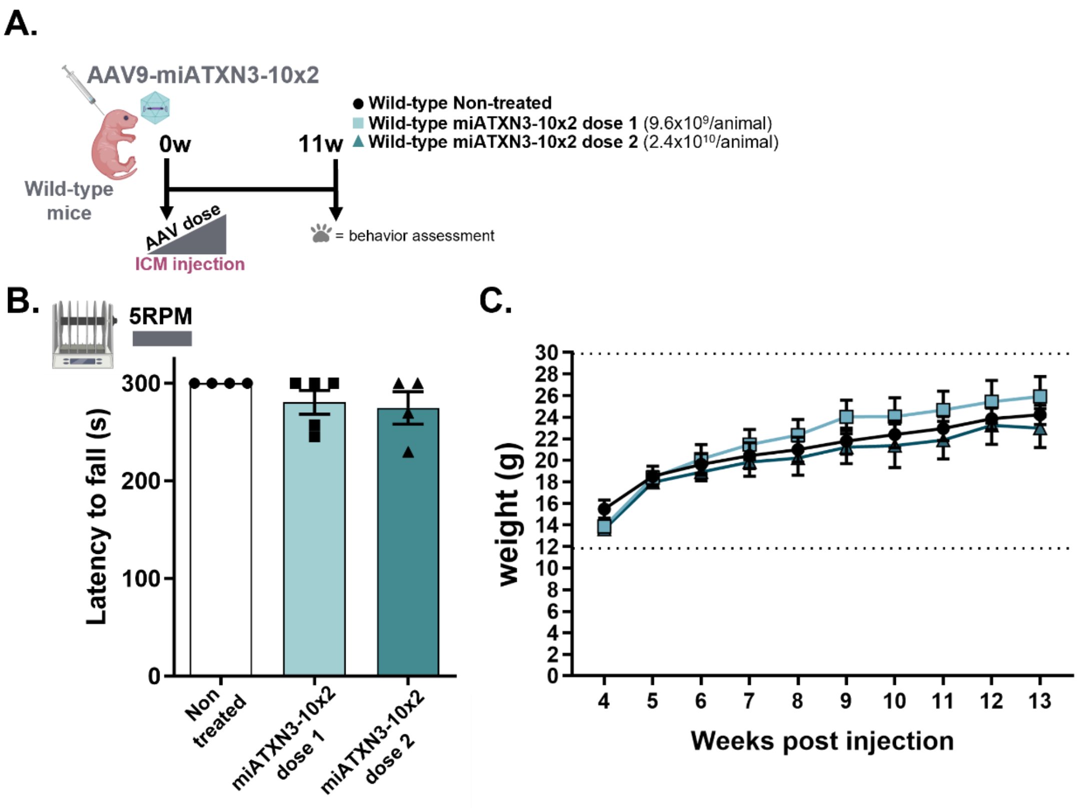


**Supplementary Figure 4: rAAV9-miATXN3-10x2 ICM administration does not affect stationary rotarod performance in wild-type mice**. **(A)** Experimental design showing wild-type mice injected with rAAV9-miATXN3-10x2 at PND1 via ICM administration. Stationary rotarod performance at a constant velocity of 5 rotations per minute (RPM) was assessed at 11 weeks post-injection. **(B)** The mean latency to fall (in seconds) is presented. No significant differences were observed between non-treated mice and those treated with rAAV9-miATXN3-10x2. Statistical analysis was performed using One-way ANOVA, followed by Dunnett's multiple comparisons post-hoc test. **(C)** The weight of wild-type animals was recorded weekly following vector administration of rAAV9-miATXN3-10x2. No significant differences in body weight were observed between experimental groups at any time point. Statistical analysis was performed using two-way ANOVA with simple effects and Tukey's multiple comparisons test. N=4-5 mice per experimental group. Data presented as mean ± SEM.

**Supplementary Table 1. Target *ATXN3* sequences and artificial miRNA antisense sequences directed against *ATXN3*-resident SNPs.**

| SEQ ID | NUCLEOTIDE SEQUENCE | DESCRIPTION |
|---|---|---|
| Target sequence at exon 8 (rs1048755 - Adenine) | 5'-ccuggaacgaauguuagaagca-3' | *ATXN3* mRNA sequence targeted by miATXN3-8 |
| Target sequence at exon 8 (rs1048755 - Guanine) | 5'-ccuggaacgaguguuagaagca-3' | *ATXN3* mRNA sequence not targeted by miATXN3-8 |
| miATXN3-8 sequence | 5'-ugcuucuaacauucguuccagg-3' | miRNA antisense sequence targeting *ATXN3* mRNA at rs1048755 (Adenine) |
| Target sequence at exon 10 (rs12895357 - Cytosine) | 5'-gcagcagcagcgggaccuauca-3' | *ATXN3* mRNA sequence targeted by miATXN3-10 |
| Target sequence at exon 10 (rs12895357 - Guanine) | 5'-gcagcagcaggggaccuauca-3' | *ATXN3* mRNA sequence not targeted by miATXN3-10 |
| miATXN3-10 sequence | *5'-ugauaggucccgcugcugcugc-3'* | miRNA antisense sequence targeting *ATXN3* mRNA at rs12895357 (Cytosine) |

Single nucleotide variants at SNP rs1048755 ($A^{669}$TG/$G^{669}$TG) and SNP rs12895357 ($C^{987}$TG/$G^{987}$TG) are underlined.

**Supplementary Table 2. Real-time quantitative PCR Primers**

| Species/Gene | Forward Primer | Reverse Primer | Ta( ºC) |
|---|---|---|---|
| Human *ATXN3** | TCCAACAGATGCATCGACCA | ACATTCGTTCCAGGTCTGTT | 56 |
| Human *ATXN3*** | TCTAGGTAAGGCCTGCTCAC | GCAAAAATCACATGGAGCTCGTA | 56 |
| Mouse *Atxn3* | GCAGATGATCAAGGTCCAACAG | TGAGGGCACTCTGCTCTTTC | 56 |
| Mouse *Hprt* | CTTCCTCCTCAGACCGCTTT | TCATCGCTAATCACGACGCT | 56 |
| Mouse *Aif1* | CTGGAGGGGATCAACAAGCAAT | AAGGCTTCAAGTTTGGACGG | 54 |
| Mouse *Gfap* | GGCTGCGTATAGACAGGAGG | CCTCCTCCAGCGATTCAACC | 54 |

Ta – Annealing temperature. *Primer sequence used for human *ATXN3* mRNA detection in samples from the SCA3 YAC84Q mouse model. **Primer sequence used for human *ATXN3* mRNA detection in samples from the SCA3 69Q mouse model.
